# Gag proteolysis drives HIV-1 envelope conformational closure through the gp41 cytoplasmic tail

**DOI:** 10.64898/2026.10.02.756164

**Authors:** David J. Williamson, Cecilia Zaza, Irene Carlon-Andres, Harry Holmes, Alessia Gentili, Joseph W. Thrush, James L. Daly, Michael Keegan, Katherine J. Doores, Tomas Malinauskas, Stuart Neil, Michael H. Malim, Sabrina Simoncelli, Sergi Padilla-Parra

## Abstract

HIV-1 must present a fusion-competent envelope glycoprotein (Env) to enter host cells while concealing the very conformation targeted by broadly neutralising antibodies (bnAbs). How the virus resolves this conflict has remained unclear. Using a minimally invasive nanobody-labelling strategy targeting Env’s V4 loop, we combine single-virus FRET, fluorescence lifetime cross-correlation spectroscopy, and DNA-PAINT super-resolution microscopy to resolve Env conformation on individual, intact virions from the primary isolate JR-FL. We show that immature virions predominantly display an open Env conformation, while Gag maturation drives acquisition of the tightly closed, fusion-competent State 1 that harbours the principal bnAb epitope. Truncating the Env cytoplasmic tail abolishes this regulation, identifying the gp41 tail as the conduit linking Gag lattice maturation to Env’s conformational fate. We further show that bnAb b12 preferentially binds open Env on immature virions, supporting a decoy mechanism that shields mature virions from immune recognition, with direct implications for HIV-1 vaccine design.

## INTRODUCTION

The envelope glycoprotein (Env) of HIV-1 is a trimeric Class I viral fusogen that mediates host-cell entry and is the principal target of broadly neutralising antibodies^1^. Env is intrinsically dynamic, transitioning between closed, prefusion-competent conformation (State 1) and more open conformations that determine both receptor engagement and antibody susceptibility^1,2^. The closed State 1 conformation, first inferred from single-molecule FRET efficiency values on intact virions by Munro et al.,^1^ represents the state targeted by the most potent broadly neutralising antibodies. However, State 1 has never been directly visualised or independently corroborated by an orthogonal, distance-resolving structural method on intact native virions; it has only ever been inferred spectroscopically from FRET efficiency values. By contrast, structures corresponding to the more open smFRET-defined states have been solved using stabilised, soluble SOSIP ectodomains engaged with CD4 and/or antibody ligands^2^. Whether and when the elusive State 1 conformation is physically acquired during the viral life cycle, and what geometry it adopts, has therefore remained unknown.

HIV-1 maturation is driven by proteolytic cleavage of the Gag polyprotein^3^ and is essential for infectivity. Previous studies have suggested functional coupling between Gag processing and Env conformation^4–6^, but the molecular mechanisms linking lattice rearrangement to Env conformational state at the single-particle level have not been resolved. A key limitation has been the inability of existing structural approaches to access native Env conformations on intact virions: cryo-EM structures of Env, including recent maps of CD4-engaged intermediates and expanded conformational landscapes, rely on soluble SOSIP ectodomains carrying stabilisation mutations that preclude adoption of the truly native closed state^7^. Cryo-electron tomography has begun to access native, membrane-embedded Env directly, resolving trimer positioning and structural heterogeneity on the Gag lattice and, in complex with CD4, intermediate open states at membrane-membrane interfaces^8^; however, a maturation-driven transition into the closed state on native virions has not been resolved by this approach, even in a dedicated recent study^9^. Together with the absence of any structure corresponding to the smFRET-defined State 1, this leaves open whether the native closed conformation is structurally distinct from existing SOSIP- or cryo-ET-derived approximations, and whether its acquisition is coupled to Gag maturation specifically, rather than to receptor engagement alone.

Here, we combine a minimally invasive AlphaFold2-guided^10^ ALFA-tag labelling strategy^11^ with fluorescence lifetime cross-correlation spectroscopy^12^ (FLCCS), DNA-PAINT^13^ super-resolution microscopy and single-particle FRET^6^ to resolve Env conformational states, including, for the first time, direct visualisation of the closed State 1 conformation by DNA-PAINT, on individual HIV-1 JR-FL^14^ immature and mature virions. We show that immature virions predominantly carry open Env trimers, while maturation drives a population shift toward a native closed conformation more compact than any previously proposed structural approximation, with V4-V4 distances of ∼ 7 nm, as compared to the SOSIP-anchored closed structure with V4-V4 distances of around 9.4 nm (i.e., PBD 5T3X). The gp41 cytoplasmic tail is essential for this regulation, identifying it as the structural conduit through which Gag lattice remodelling is communicated to the Env trimer. Together, these findings establish that the native, neutralisation-sensitive closed Env conformation is acquired only upon maturation, with direct implications for vaccine strategies targeting native Env structure.

## RESULTS

### AlphaFold2-guided ALFA-tag insertion enables minimally invasive Env labelling on intact virions

Accurate labelling of viral Env proteins without perturbing receptor binding or fusion activity is a prerequisite for studying their conformational dynamics on intact virions via fluorescence spectroscopy and microscopy. We selected the fourth variable (V4) loop of the primary CCR5-tropic isolate JR-FL^14^ as an insertion site because it is highly surface-exposed and functionally tolerant (Figure 1A, B). AlphaFold2^10^ was used to screen candidate positions for ALFA-tag insertion across the V4 loop (Table S1). Structural models (*n* = 26) were evaluated for tag conformation and nanobody accessibility, requiring that the ALFA-tag adopt an outward-facing α-helical conformation compatible with the anti-ALFA nanobody (nbALFA) binding pocket (PDB: 6I2G) (Figure 1C; S1–S5). One favourable insertion site (UIDs 415d2) and one unfavourable control (UID 637e9) were selected for experimental validation. Both V4-ALFA-tagged Env variants were expressed, properly cleaved into gp120 and gp41 subunits, and efficiently incorporated into HIV-1 particles. Single-cycle infectivity assays in TZM-bl reporter cells confirmed that ALFA-tag insertion did not impair viral entry, and nbALFA binding did not inhibit infectivity (Figure 1D; S6). Cell–cell fusion assays corroborated these findings (Figure S7). To validate specific Env labelling on intact particles, we performed fluorescence lifetime cross-correlation spectroscopy (FLCCS)^15^ on virions co-labelled with Gag-EGFP, reporting virions, and nbALFA-ATTO594, reporting Env engagement (Figure 1E). Robust cross-correlation was observed for the 415d2 construct (Figure 1F), with no detectable interaction for the structurally disfavoured 637e9 variant (Figure 1G), directly validating the AlphaFold2 accessibility predictions. Together, these results establish 415d2 as a fully functional, specifically labelled Env construct and validate AlphaFold2-guided screening as an efficient strategy for epitope tag insertion into viral glycoproteins.

**Figure 1.**
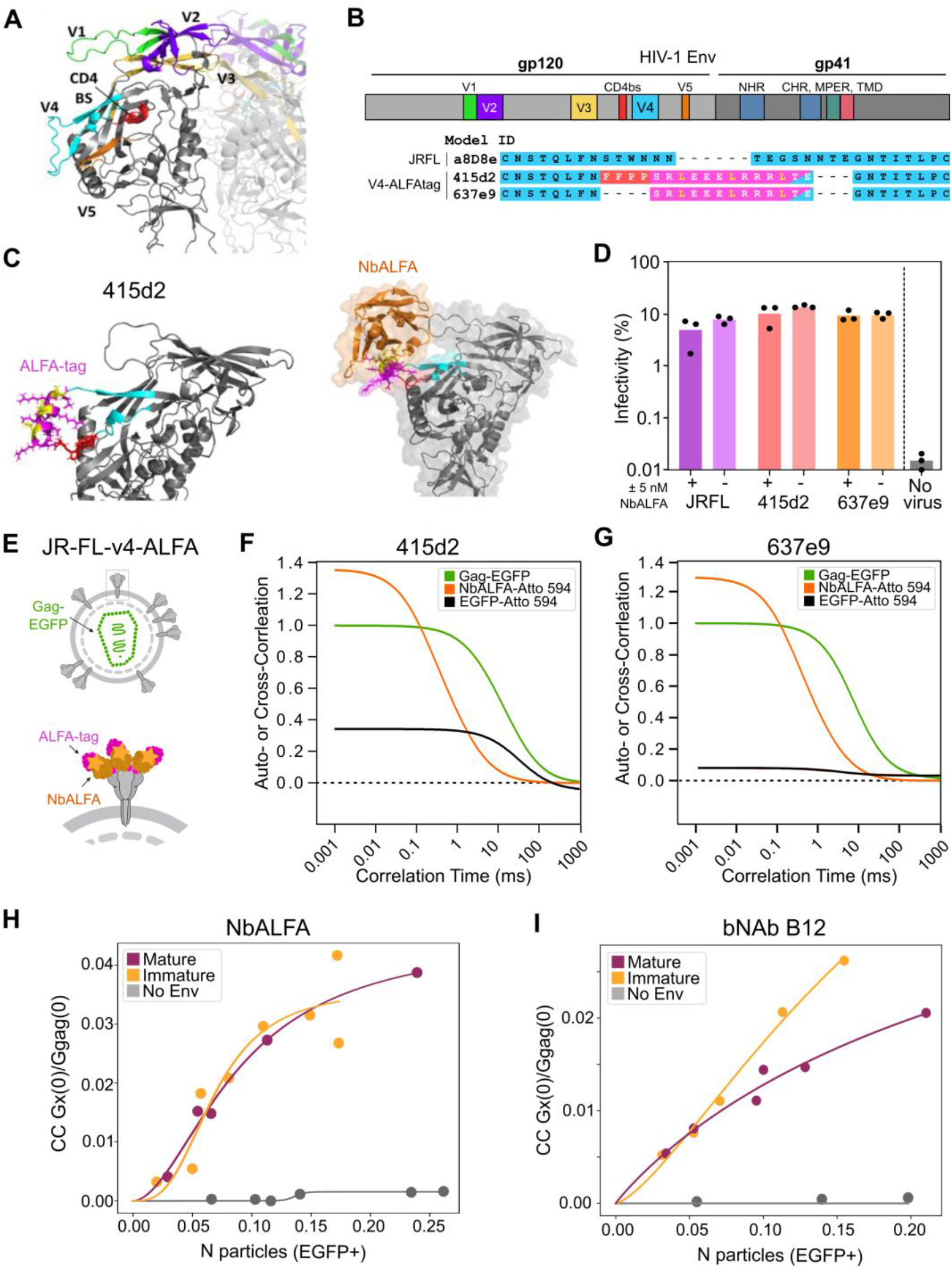
AlphaFold2-guided ALFA-tag insertion enables specific Env labelling on intact HIV-1 virions and reveals preferential binding of broadly neutralising antibody b12 to open Env conformations on immature particles. **(A)** Structural model of the JR-FL Env gp120 subunit, showing the variable loops (V1–V5, coloured) and the CD4 binding site (CD4bs). **(B)** Domain organisation of HIV-1 Env (gp120 and gp41 subunits, top) and sequence alignments of the V4 loop region for wild-type JR-FL and the two-candidate ALFA-tag insertion sites (415d2 and 637e9); inserted ALFA-tag residues are highlighted in magenta. **(C)** AlphaFold2 models of the 415d2 insertion design. Left, predicted V4 loop structure with the ALFA-tag (magenta) adopting an outward-facing α-helical conformation; interface leucines (yellow); linker residues (red). Right, docking model of nbALFA (orange) onto the predicted tag conformation. V4 loop, cyan; ALFA-tag, magenta; interface leucines, yellow; linker residues, red; nbALFA, orange. **(D)** Single-cycle infectivity of pseudotyped HIV-1 particles bearing JR-FL, 415d2, or 637e9 Env, in the presence or absence of 5 nM nbALFA, normalised to untreated JR-FL. **(E)** Schematic of the FLCCS assay: intact JR-FL-V4-ALFA virions are co-labelled with Gag-EGFP (virion reporter) and nbALFA–ATTO594 (Env reporter); cross-correlation between the two channels reports co-diffusion of labelled Env on intact virions. **(F,G)** Representative fluorescence autocorrelation and cross-correlation curves for virions bearing the structurally favourable 415d2 **(F)** or unfavourable 637e9 **(G)** insertion constructs. Robust cross-correlation between Gag-EGFP and nbALFA–ATTO594 was exclusively observed for 415d2, confirming tag accessibility on intact virions and validating the AlphaFold2 predictions. **(H,I)** FLCCS binding curves for mature (purple) and immature (orange) JR-FL-v4-ALFA virions, with no-Env particles as negative control, as a function of virion concentration (GFP+ particles). **(H)** NbALFA–ATTO594 binding to ALFA-tagged Env showed comparable labelling of mature and immature particles. **(I)** Binding of the conformation-sensitive broadly neutralising antibody b12 is markedly increased on immature relative to mature virions, with higher maximum binding capacity, lower apparent dissociation constant, and positive cooperativity, indicating a greater abundance of open Env conformations on immature particles.

### Immature HIV-1 virions preferentially expose open Env conformations

Having established specific Env labelling, we next asked whether HIV-1 Env conformational accessibility differs between immature and mature virions at the population level using conformation-sensitive antibody binding quantified by FLCCS with pulsed interleaved excitation (PIE)^16^, which alternates the excitation lasers so that photons can be assigned to their source before cross-correlation, suppressing spectral bleed-through between channels^17^. As a prerequisite, we confirmed that Env surface exposure is equivalent between immature and mature JR-FL-V4-ALFA (415d2) virions using nbALFA-ATTO594, which binds the ALFA tag independently of Env conformation (Figure 1H). Both populations yielded indistinguishable binding parameters across a range of viral concentrations (B*max* = 0.03, K*d* = 0.09, Hill coefficient *h* = 1.5) — K*d* is expressed here, and throughout, in units of particle concentration rather than molar ligand concentration, since nanobody or antibody was held in molar excess while virion number was titrated —, ruling out differences in Env density as a confound for subsequent experiments. We then probed conformational accessibility using the broadly neutralising antibody (bnAb) b12^18^, which selectively recognises the open, non-CD4-bound conformation of Env by engaging an epitope overlapping the CD4 binding site^8^, making its binding a direct readout of open-state accessibility. Among a panel of bnAbs spanning the major Env epitope classes, b12 was among the most potent against JR-FL-V4-ALFA virions and the most potent of the CD4-binding-site-directed antibodies tested, supporting its selection for these conformation-sensitive binding assays (Figure S8).

The binding curves for mature and immature virions were strikingly different when increasing the number of particles (Figure 1I): immature virions showed a substantially higher maximum binding capacity (B*max* = 0.06 vs. 0.04), a lower apparent dissociation constant (K*d* = 0.21 vs. 0.30), and positive cooperativity (Hill coefficient *h* = 1.41), whereas mature virions displayed near non-cooperative binding (*h* = 0.94). The steeper, higher-amplitude binding curve on immature particles indicates a greater abundance of b12-accessible, open Env conformations relative to mature virions, providing the first direct population-level evidence that HIV-1 maturation shifts the Env conformational equilibrium away from the open state.

### Maturation shifts the Env conformational equilibrium toward a compact native closed state

To directly resolve Env conformational states at the level of individual trimers on intact virions, we applied DNA-PAINT super resolution imaging to JR-FL-V4-ALFA (415d2) particles (Figure 2A-B), achieving lateral localisation precisions of 2.0–2.5 nm (Cramér–Rao lower bound; Figure S9), corresponding to an effective spatial resolution of ∼5–6 nm, surpassing previous fluorescence-based imaging of Env in HIV-1 viruses^4,19^.

**Figure 2.**
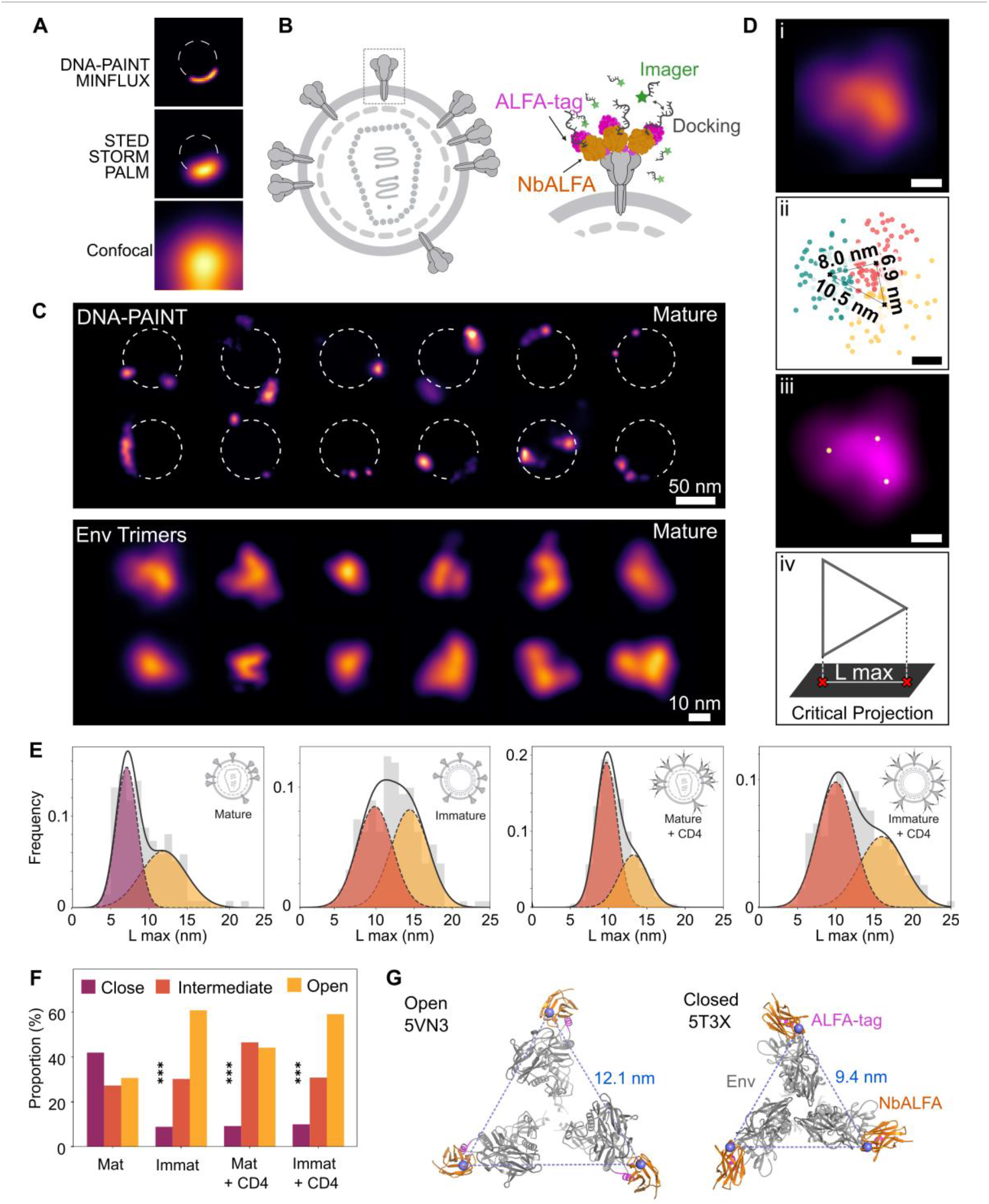
DNA-PAINT super-resolution imaging resolves a native compact closed Env conformation on mature HIV-1 virions. **(A)** Schematic illustrating the resolution capabilities of different fluorescence microscopy modalities for visualising Env clusters on intact viral particles. **(B)** DNA-PAINT labelling strategy: nbALFA bearing a DNA docking strand is bound to the ALFA-tagged Env trimer and transiently engaged by fluorescent imager strands. **(C)** Representative 2D DNA-PAINT images acquired under TIRF illumination of individual mature HIV-1 virions (dashed circles, top panel; scale bar, 50 nm); magnified views of individual Env trimers resolved on the virion surface (bottom panel; scale bar, 10 nm). **(D)** Analysis pipeline for single-trimer inter-protomer distance measurements: (i) representative DNA-PAINT image of a single Env trimer; (ii) three-component Gaussian Mixture Model (GMM) fit to single-molecule localisations, yielding protomer centre-of-mass positions and pairwise distances; (iii) overlay of GMM-fitted protomer positions (white dots) on the trimer image; (iv) schematic illustrating extraction of the maximal side length (L max) from the critical projection to mitigate 3D-to-2D projection artefacts. **(E)** Distributions of L max for mature (top left) and immature (top right) virions, and for mature (bottom left) and immature (bottom right) virions treated with soluble CD4, fitted with a free two-component Gaussian model (closed, purple; intermediate, orange; open, yellow). Mature virions display a predominant closed population (∼7 nm) largely absent from immature virions; sCD4 treatment drives mature trimers toward intermediate and open states while immature virions are largely refractory. **(F)** Quantification of the fractional areas of closed (< 8 nm), intermediate (8–11 nm), and open (> 11 nm) trimer populations across the four conditions (mature, immature, mature + CD4, immature + CD4). Data are mean ± s.e.m. Statistical comparisons shown are for the closed conformation only (***p < 0.001); full pairwise statistics for all conformational populations are provided in Supplementary Table 2. **(G)** AlphaFold2 models of the JR-FL-v4-ALFA:nbALFA complex docked onto cryo-EM reference structures for the open (PDB: 5VN3, top) and closed (PDB: 5T3X, bottom) and Env conformations, with predicted Val122–Val122 inter-nanobody distances of 9.4 nm and 12.1 nm, respectively. The SOSIP-stabilised ectodomain structure 5T3X, conventionally referred to as closed, yields a predicted inter-nanobody distance exceeding our experimentally measured native closed-state distance of ∼7 nm, suggesting it corresponds more closely to an intermediate conformation in our nomenclature. The experimentally measured closed-state distance of ∼7 nm is more compact than the 5T3X-anchored prediction, consistent with 5T3X representing a SOSIP-stabilised ectodomain carrying engineered mutations that preclude adoption of the truly native closed conformation.

Representative DNA-PAINT images of intact mature HIV-1 virions revealed individual Env complexes distributed across the virion surface (Figure 2C, top panel), with magnified views resolving the characteristic triangular arrangement of gp120 subunits within individual trimers (Figure 2C, bottom panel and S10). A three-component Gaussian Mixture Model (GMM) was fitted to the single-molecule localisations^20^ within each trimer to determine protomer centre-of-mass positions and inter-subunit distances (Figure 2D, i-iii). Because Env sits on the curved viral membrane while DNA-PAINT captures only a 2D projection, foreshortening can compress the shorter edges of an imaged trimer; we therefore only analysed the maximal side length (L max)^21^ of each trimer (Figure 2D, iv), a choice validated by simulating 3D equilateral trimers of known inter-protomer spacing (7, 10 and 12 nm) projected into 2D under matched imaging conditions, which confirmed that L max recovers the true distance with high fidelity ((7.3 ± 0.9), (9.9 ± 0.9) and (11.8 ± 0.7) nm, respectively; Figure S11).

To classify individual trimers, we examined the pooled L max distributions for each condition (Figure 2E). Mature virions were unique in showing a distinct low-distance population centred below 8 nm; every other condition (i.e., immature virions and both CD4-treated populations) lacked this population entirely, with their shortest distances instead clustering around 10 nm (Table S2). We used this separation to define three classes: closed (L max < 8 nm), intermediate (8–11 nm), and open (>11 nm) (Figure 2F).

Applying this classification revealed striking maturation-dependent differences (Figure 2F). Mature virions (*N* = 137 trimers) showed ∼42% closed, ∼27% intermediate , and ∼31% open trimers. In contrast, immature virions (*N* = 191 trimers) shifted the distribution markedly toward the open state, with ∼61% of trimers classified as open, ∼30% intermediate, and only ∼9% closed (Figure 2F). Addition of soluble CD4 (sCD4) to mature virions (*N* = 211 trimers) redistributed trimers toward intermediate and open states (∼47% intermediate, ∼44% open, ∼9% closed), while immature virions (*N* = 395 trimers) were largely insensitive to sCD4 (∼31% intermediate, ∼59% open, ∼10% closed), consistent with their already-open baseline conformation. These data provide a nanoscale structural basis for the differential b12 binding observed by FLCCS (Figure 1I).To independently validate these conformational assignments, we generated AlphaFold2 models of the JR-FL-v4-ALFA:nbALFA complex superposed onto cryo-EM reference structures for the closed (PDB: 5T3X)^22^ and open (PDB: 5VN3)^18^ conformations, measuring Val122-Val122 distances between nanobody pairs, corresponding to the DNA docking strand attachment points and thus directly comparable to our DNA-PAINT measurements (Figure 2G). The predicted open distance of 12.1 nm agrees closely with our experimental measurement of 12 ± 2 nm. For the closed state, however, the AlphaFold prediction of 9.4 nm anchored to 5T3X exceeds our experimental measurement of 7 ± 1 nm; a discrepancy we attribute to 5T3X being a soluble SOSIP ectodomain stabilised by engineered mutations, antibody-bound, and lacking the cytoplasmic tail essential for native conformational regulation. We propose that the 7 nm conformation observed predominantly on mature virions represents the truly native closed conformation, more compact than any previously captured by structural approaches relying on stabilised ectodomains.

### The gp41 cytoplasmic tail transduces Gag maturation signals to regulate Env conformation

To probe the mechanistic basis of maturation-dependent Env regulation, we examined virions carrying a C-terminal truncation of Env (EnvΔCT)^23^, known to enhance Env incorporation30. Both FLCCS (Figure S12) and quantitative DNA-PAINT (qPAINT)^24^, which infers protein copy number from docking-strand and imager-strand binding kinetics, confirmed elevated Env incorporation in both immature and mature EnvΔCT particles (∼3-fold increase by DNA-PAINT; p-value < 0.0001; Figure 3), with indistinguishable b12 binding parameters between maturation states (B*max* = 59.6, K*d* = 9.6, *h* = 1.4 for Mature ΔCT and B*max* = 145.6, K*d* = 10.4, *h* = 1.6 for Immature ΔCT; Figure 4A). This indicates that cytoplasmic tail truncation specifically abolished the maturation-dependent difference in Env conformational accessibility.

**Figure 3.**
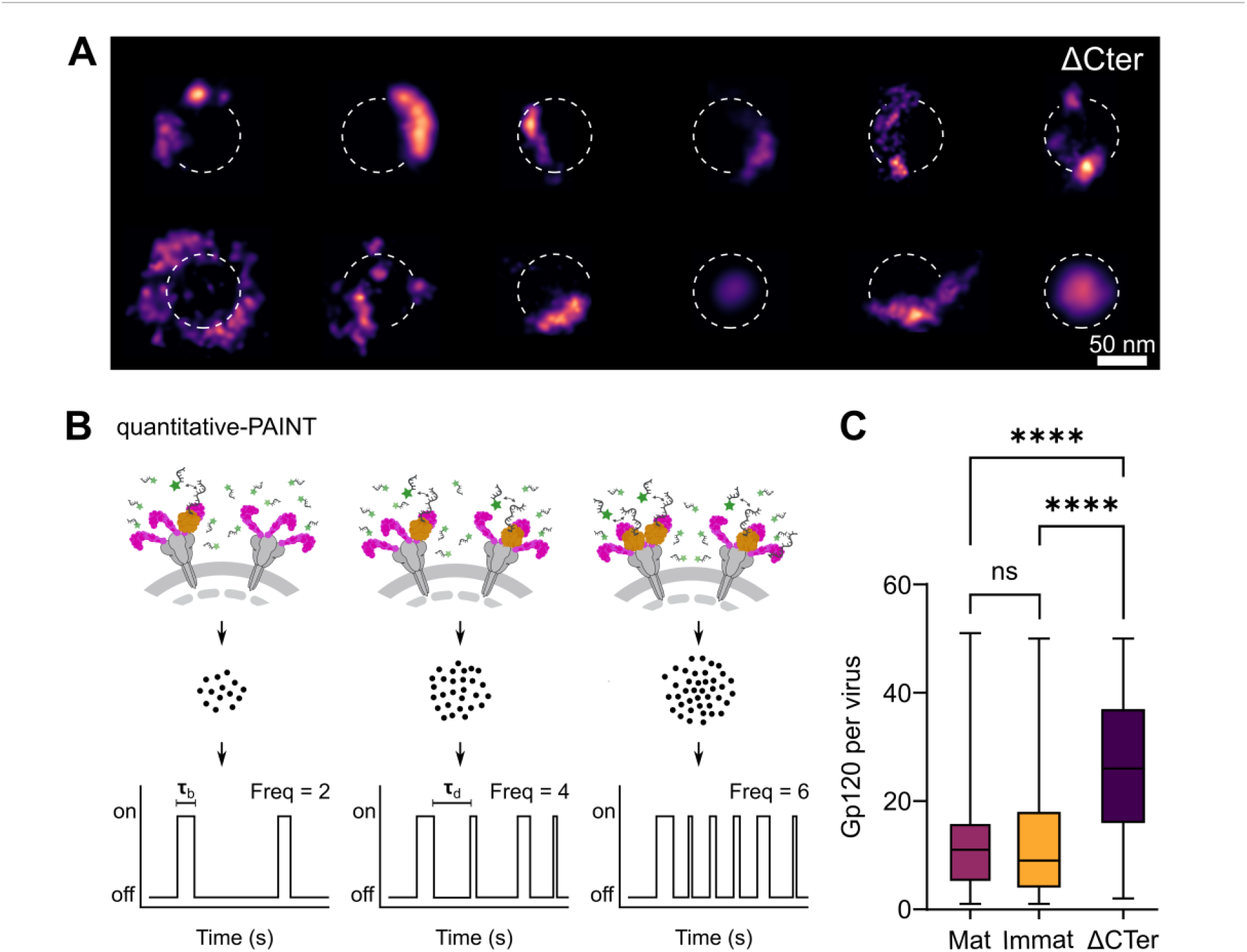
qPAINT quantification of Env incorporation per HIV-1 particle. **(A)** Representative 2D DNA-PAINT images of individual HIV-1 virions bearing JR-FL Env V4-ALFA with a truncated gp41 cytoplasmic tail (ΔCT). Dashed circles indicate virion boundaries. Scale bar, 50 nm. **(B)** Schematic of the quantitative DNA-PAINT (qPAINT) approach used to determine gp120 copy number per virion. Fluorescently labelled imager strands transiently bind complementary docking strands conjugated to nbALFA targeting V4-ALFA labelled Env. The resulting fluorescence intensity traces exhibit characteristic binding on- and off-times. Using the binding frequency of a single imager–docking strand pair as a reference, the number of docking sites within each localisation cluster is calculated from the ratio of the observed cluster binding frequency to that of a single site. **(C)** gp120 copy number per virion determined by qPAINT for mature and immature JR-FL V4-ALFA particles and mature ΔCT V4-ALFA particles. ΔCT virions exhibit approximately threefold higher Env incorporation than wild-type particles (*P* < 0.0001), consistent with increased Env surface density following cytoplasmic tail truncation.

**Figure 4.**
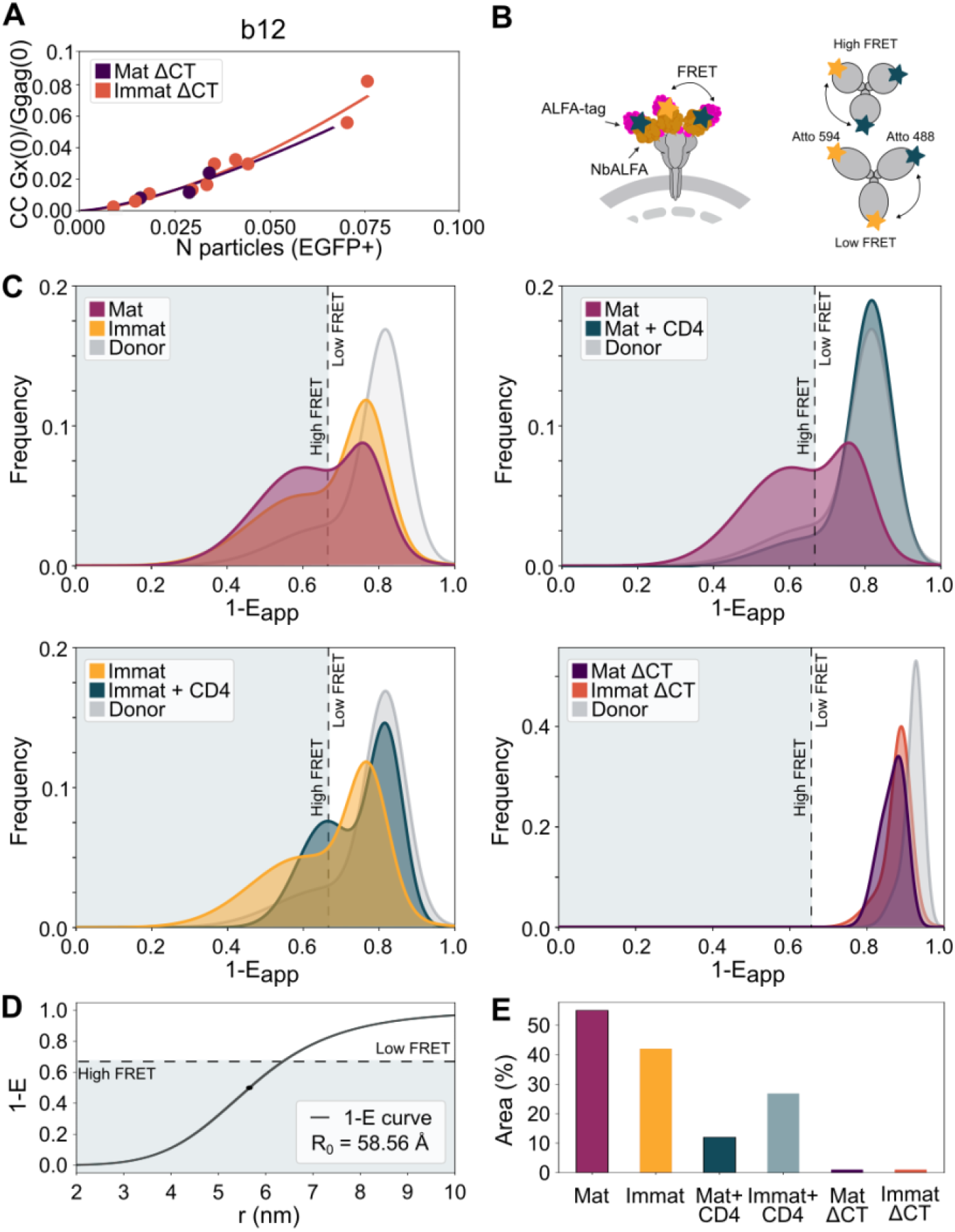
Single-virion FRET confirms maturation-dependent acquisition of the closed Env conformation and identifies the gp41 cytoplasmic tail as essential for its regulation. **(A)** FLCCS binding curves for b12 on mature (dark purple) and immature (orange) EnvΔCT virions, showing indistinguishable binding parameters between maturation states and markedly elevated maximum binding capacity relative to wild-type virions (Fig. 1I), confirming increased Env incorporation on ΔCT particles. The high Env surface density on EnvΔCT virions precluded reliable isolation of individual trimers by DNA-PAINT, necessitating single-virion FRET to probe conformational states in this background. **(B)** Schematic of the single-virion FRET assay: intact JR-FL-v4-ALFA virions are stochastically co-labelled with nbALFA–ATTO488 (donor) and nbALFA– ATTO594 (acceptor); closed trimers bring donor–acceptor pairs into proximity, yielding high FRET, while open trimers produce low FRET. **(C)** Normalised 1-Eapp distributions for the indicated conditions, with donor-only controls shown in grey and a dashed line at 1-Eapp = 0.67 demarcating high-FRET (closed, left) and low-FRET (open, right) populations. Mature virions display a prominent high-FRET population relative to immature virions (top left); sCD4 treatment drives mature trimers toward low FRET while immature virions are partially refractory (top right and bottom left); both MatΔCT and ImmatΔCT virions show distributions indistinguishable from the donor-only control, indicating a complete failure to acquire stable closed conformations regardless of maturation state (bottom right). **(D)** Theoretical 1-E versus inter-dye distance curve calculated using the experimentally determined Förster radius R₀ = 58.56 Å, with the high-FRET/low-FRET threshold indicated. **(E)** Quantification of the fractional area in the high-FRET region across all six conditions (Mat, Immat, Mat+CD4, Immat+CD4, MatΔCT, ImmatΔCT). Data are mean ± s.e.m.

The high Env density on EnvΔCT particles precluded reliable isolation of individual trimers by DNA-PAINT, so we turned to single-virion FRET to directly probe conformational states of JR-FL-V4-ALFA viral particles, stochastically labelled with nbALFA–ATTO488 (donor) and nbALFA–ATTO594 (acceptor) (Figure 4B and S13). These experiments yielded apparent FRET efficiency distributions (1-Eapp), drawn from more than 2,000 individual particles per condition, fully consistent with DNA-PAINT conformational assignments (Figure 4C). Mature virions exhibited a prominent high-FRET population extending well below the 1-Eapp threshold of 0.67 (Figure 4D), clearly distinct from the donor-only control, which peaks at ∼ 0.8 with negligible signal in this region, directly reporting a substantial closed Env population. This directly reports a substantial closed Env population. The threshold was defined as the donor-only μ + 2σ cutoff from a Gaussian mixture model fit to the donor-only control distribution. Immature virions, on the other hand, displayed a distribution shifted toward donor-only levels, with markedly less area below the threshold. Quantification confirmed that maturation shifts the conformational equilibrium toward the closed state (∼55% mature vs ∼40% immature; Figure 4E), fully consistent with DNA-PAINT. Addition of sCD4 drove near-complete collapse of the closed population in mature virions, while immature virions retained residual signal above donor-only, in keeping with the DNA-PAINT observation that immature trimers are largely refractory to sCD4-induced conformational change (Figure 4E).

Strikingly, both immature and mature EnvΔCT virions displayed 1-Eapp distributions indistinguishable from donor-only controls (Figure 4D), indicating a complete failure to achieve stable closed conformations regardless of Gag processing status. These findings suggest that the gp41 cytoplasmic tail is required for maturation-dependent conformational regulation, and that its interaction with the underlying matrix lattice allosterically governs the acquisition of the closed State 1 conformation.

## DISCUSSION

Our findings establish viral maturation as the primary determinant of HIV-1 Env conformational state on intact native virions, resolving a longstanding question in HIV-1 biology. While Munro et al. first identified the closed State 1 conformation by single-molecule FRET^1,7,25^, the relationship between maturation^26^ and conformational state was unknown. We now show that this conformation is specifically acquired upon Gag cleavage, revealing a direct mechanistic link between the structural reorganisation of the viral interior and the conformational landscape of the exterior glycoprotein (Figure 5).

**Figure 5.**
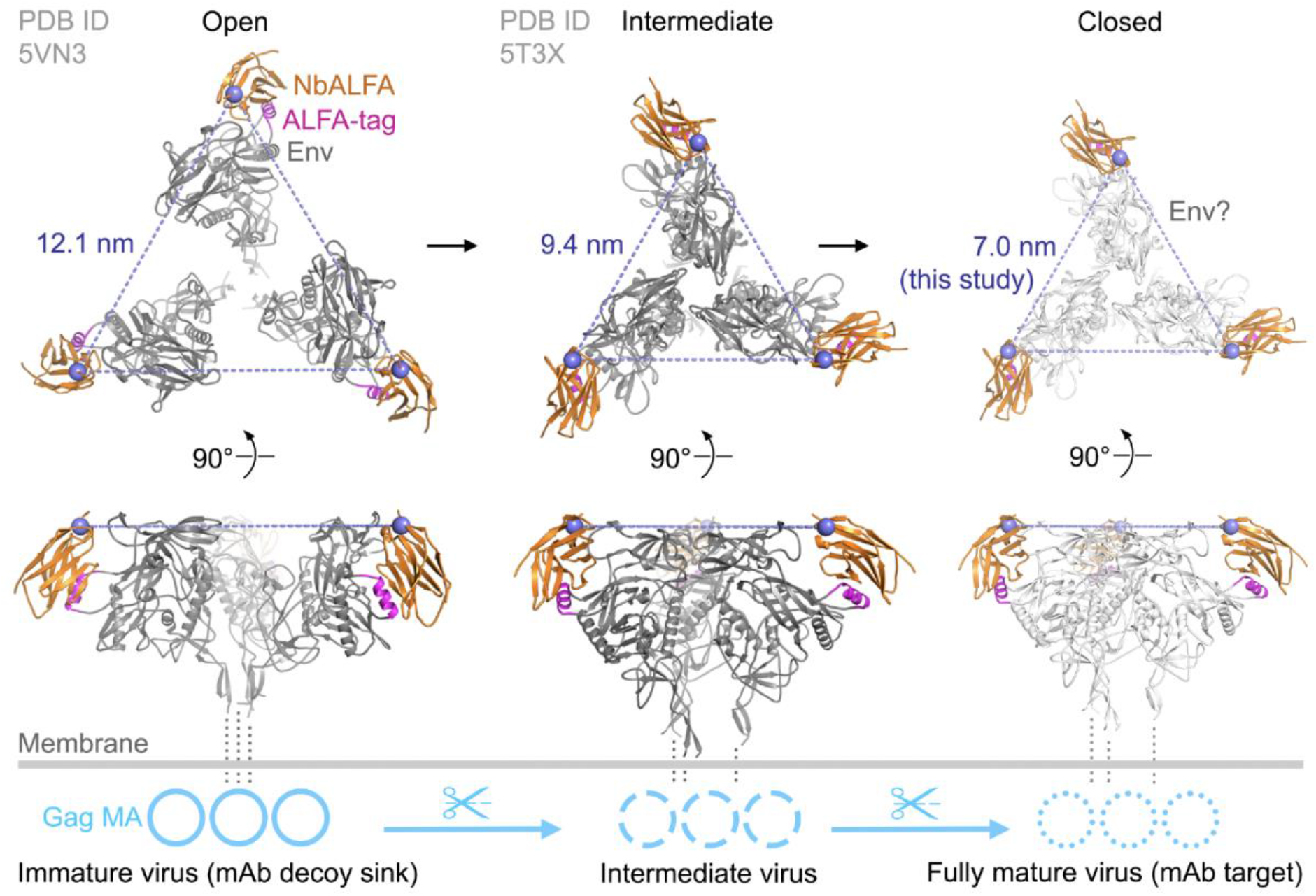
Cartoon summarising the model for the correlation between Env conformational dynamics and maturation. Immature viruses predominantly present a fully open JR-FL Env conformation (left-hand side), corresponding to V4-V4 distances of ∼ 12nm (e.g. PBD 5VN3). Upon maturation and Gag processing, JR-FL Env redistributes toward a fully closed conformation (right-hand side), with unknown atomic structure and V4-V4 distances of ∼ 7 nm. The ALFA-tag nanobody (orange) and the fluorophore (blue sphere) are also represented, as DNA-PAINT directly measures distances between the chromophores rather than between the Env protomers themselves. An intermediate, partially closed structure was also found, with V4-V4 distances of ∼ 9 nm (PBD 5T3X). This structure does not represent the fully closed, native metastable Env conformation but rather a partially open Env state

Using three complementary single-molecule and super-resolution imaging approaches—FLCCS, DNA-PAINT and FRET—applied to intact, infectious JR-FL virions labelled with the ALFA-tag platform, we demonstrate that the metastable closed State 1 Env conformation is specifically acquired upon maturation and depends on the integrity of the gp41 cytoplasmic tail^27^. In immature virions, Env trimers predominantly sample open and intermediate conformations, with only a minor fraction adopting the closed state. Upon Gag cleavage and maturation, this equilibrium shifts dramatically, with the largest fraction of trimers stabilised in the closed conformation (Figure 2E and 4C). This maturation-dependent shift is consistent with structural rearrangements in the matrix (MA) lattice that relieve constraints on the gp41 cytoplasmic tail^2^, enabling allosteric stabilisation of the closed ectodomain^28^. The role of the gp41 cytoplasmic tail as an allosteric regulator of Env conformation is reinforced by the ΔCT phenotype and has been suggested by others^29^. Truncation of the cytoplasmic tail abolishes maturation-dependent conformational regulation entirely^30^, producing virions with elevated Env incorporation but constitutively open trimers regardless of Gag processing status. This finding extends previous demonstration that Env clustering on the virion surface is also regulated by cytoplasmic tail–matrix interactions4,5, revealing that the cytoplasmic tail plays a dual role: organising Env spatial distribution^5^ and constraining its conformational landscape. The conserved SP2 spacer peptide, whose removal during maturation is now known to trigger MA lattice reorganisation^28^, may represent a key molecular switch in this process with its cleavage potentially relieving steric constraints on the cytoplasmic tail that otherwise prevent adoption of the closed ectodomain conformation.

The preferential binding of the broadly neutralising antibody b12 to immature virions, with greater capacity and positive cooperativity compared to mature particles (Figure 1I), raises an intriguing possibility. If open Env conformations on immature virions preferentially engage b12 and related antibodies^31^ targeting the CD4 binding site, these non-infectious particles could act as conformational decoys, absorbing neutralising antibody responses directed at open Env epitopes while the infectious mature virions present the closed conformation that is less accessible to the same antibodies^32^. This hypothesis, if confirmed, would represent a previously unrecognised immune evasion strategy operating at the level of Env conformational dynamics rather than sequence variation or glycan shielding^33^ alone.

These findings carry direct implications for HIV-1 vaccine design. SOSIP trimers^34^, the dominant current immunogen platform, are engineered to mimic the native virion-associated trimer^35^, yet our data suggest they present an engineered approximation of the closed state rather than the truly native conformation on intact mature virions. This structural discrepancy may contribute to their failure to consistently elicit broadly neutralising antibody responses^36^. Vaccine platforms that more accurately reproduce the native closed conformation, whether through intact mature virion-like particles or stabilised constructs constrained to the 7 nm inter-protomer geometry, may more faithfully present the antigenic surface targeted by the most potent broadly neutralising antibodies. Equally important is the temporal dimension: the native closed conformation exists only on mature infectious virions (Figure 2 and 4), defining a narrow window during which the immune system encounters the relevant target. Future studies should examine whether this conformational window influences the quality and breadth of the humoral immune response.

## ACKNOWLEDGEMNTS

This work was funded by the Chan Zuckerberg Initiative (2023-321188) to S.S. and S.P.-P.; the Human Frontier Science Program (LT0025/2023-C) to C.Z.; the Royal Society Dorothy Hodgkin Fellowship (DHF\R1\191019 and DHF\R\251006) to S.S.; the European Research Council (ERC-2019-CoG-863869 FUSION) to S.P.-P.; the Wellcome Trust Senior Research Fellowship (WT207442/Z/17/Z) and MRC (G0801937) to S.N.; and the Wellcome Trust (222433/Z/21/Z, 225128/Z/22/Z) to M.H.M. and J.D. We thank the Nikon Imaging Centre at King’s College London for support with confocal microscopy.

## AUTHOR CONTRIBUTIONS

Conceptualisation: D.J.W., S.S., S.P.-P. Methodology and validation: D.J.W., C.Z., I.C.-A., H.H., A.G. J.D., J.W.T..,M. K., K.J.D., T.M., S.N., M.H.M., S.S., S.P.-P. Formal analysis: D.J.W., C.Z., I.C-A., , H.H., A.G. J.D., J.W.T..,M. K., K.J.D., T.M., S.S. Resources: S.S., S.P.-P. Data curation: D.J.W., C.Z., S.S., S.P.-P. Writing—original draft: S.P.-P., S.S. Writing—review and editing: all authors. Visualisation: D.J.W., C.Z., T.M. S.S., S.P.-P. Supervision: S.S., S.P.-P. Project administration and funding acquisition: S.S., S.P.-P.

## MATERIAL AND METHODS

### Cell culture

All cell lines were maintained at 37°C in a humidified atmosphere with 5% CO₂ without antibiotics. Producer cell lines (Lenti-X 293T, Takara Bio, 632180) and reporter cells (TZM-bl, NIH AIDS Research & Reference Reagent Program) were propagated in phenol-red-free DMEM/F12 (Thermo, 21041033) supplemented with 10% fetal bovine serum (FBS).

### Plasmid constructs

Plasmids pR8ΔEnv and pcRev were provided by Dr Greg Melikyan (Emory University). The pCAGGS-JR-FL Env plasmid was a gift from Dr James Binley (Torrey Pines Institute for Molecular Studies). ALFA-tag insertion variants were generated by site-directed mutagenesis guided by AlphaFold2 structural models.

### Single-domain antibody labelling

Purified anti-ALFA-tag single-domain antibodies (N1505, Nano-tag) were reduced with 15 mM TCEP for 10 min on ice, buffer-exchanged into maleimide labelling buffer (100 mM potassium phosphate pH 6.4, 150 mM NaCl, 1 mM EDTA, 250 mM sucrose), and conjugated with ATTO594 or ATTO488 maleimide (12:1 molar ratio, 1.5 h on ice, pH 7.5). Unreacted dye was removed by PD-10 desalting. Degree of labelling was confirmed by SDS–PAGE and spectrophotometry.

### Virus production and concentration

Lenti-X 293T cells were seeded at 65 × 10³ cells/cm² and transfected the following day using GeneJuice (Merck, 70967; 3:1 reagent:DNA) in FluoroBrite DMEM (Gibco) with 3% FBS. Transfection mixtures contained pR8ΔEnv, pcRev, Gag–fluorescent protein, and Env at a 1:2:0.1:2 plasmid ratio. Virions were harvested 48 h post-transfection, clarified (2,000 × *g*, 5 min; 0.45 µm filtration), and concentrated by sucrose cushion ultracentrifugation (10,000 × *g*, 3–4 h, 4°C). Pellets were resuspended in PBS and stored at −80°C.

### Virus titration

Single-cycle infectivity was determined using TZM-bl reporter cells plated at 60 × 10³ cells/cm². Serially diluted virus was added 24 h after plating; cells were fixed after 48 h in 2% PFA, stained with X-Gal solution, and the proportion of β-galactosidase-positive (infected) cells quantified by brightfield microscopy.

### Western blotting

Cells and concentrated viral pellets were lysed in 1% Triton X-100 PBS with protease inhibitors. Proteins were resolved on NuPAGE 4–12% Bis-Tris gels (Life Technologies) and transferred to PVDF membranes (Millipore IPFL00010). Membranes were probed with anti-GAPDH (Proteintech), anti-p24 (24-2), anti-gp41 (10E8), or FluoTag®-X2 anti-ALFA–LICOR IRDye 680RD (NanoTag), followed by fluorescent secondary antibodies and LI-COR Odyssey Fc imaging.

### PIE–FLCCS

FLCCS measurements were performed on a MicroTime 200 (PicoQuant) at 37°C with pulsed interleaved excitation at 485 nm (Gag–eGFP) and 595 nm (sdAb–ATTO594) at 20 MHz repetition rate, focused through a 63× 1.2 NA water immersion objective (Olympus UPlanXApo). Emission was separated by a 560LP dichroic; the green channel used a 525/50 BP filter (PMA hybrid detector) and the red channel a 600LP/690-70 BP combination (SPAD). Time-correlated single-photon counting (TCSPC) was performed with a MultiHarp 150 (PicoQuant). Autocorrelation and cross-correlation curves were analysed in SymPhoTime 64 using a free 3D diffusion model.

### Single-virion FRET imaging

FRET measurements used the MicroTime 200 with pulsed excitation at 440 nm (Gag–mAmetrine) and 485 nm (nbALFA–ATTO488), through a 100× 1.2 NA oil objective and a 300 µm pinhole. Donor (525/50 BP) and sensitised acceptor (650/50 BP) emissions were detected in parallel. Apparent FRET efficiencies were extracted from >2,000 individual particles per condition using SymPhoTime 64 and custom ImageJ software incorporating automated particle picking and thresholding.

### DNA-PAINT sample preparation

Glass bottom 6 channel Ibidi µ-slides (IB-80607) were cleaned with KOH 1 M for 2 h at room temperature and washed with copious volumes of clean water. The glass surface was then air-dried and exposed to UV light for 30 minutes. Viral particles were diluted 1:10 in PBS with 2% PFA and fixed at room temperature for ten minutes, then further diluted 1:10 in PBS. Fixed viral particles were added to the glass surface for 30 minutes and then the slide was washed 3 × 5 minutes with PBS, following 30 min blocking with PBS containing 2% BSA, 0.2% Fish Skin Gelatin. 25 nM DNA coupled anti-ALFA-tag nanobody (MASSIVE-TAG-Q-ANTI-alfa with R1: 5’-TTTCCTCCTCCTCCTCCTCCT-3’) in BSA 5% in PBS for 1 hour at room temperature. Samples were washed 3 × 5 minutes with PBS before the addition of 90 nm gold nanoparticles (G-90-100, CytoDiagnostics) for 5 minutes and a final 3 × 5 minutes with PBS. For imaging, DNA imager solution (R1: AGGAGGA) with 3’ attached Cy3b fluorophore in C+ buffer (1× PBS, 1 mM EDTA (Invitrogen), 500 mM NaCl (Sigma) pH 7.4, 0.02% Tween (Sigma)) was used. For 2D-DNA-PAINT-TIRF imaging a concentration of 2.5 nM of imager was used and for DNA-PAINT MINFLUX 1 nM imager concentration was used.

### DNA-PAINT super-resolution imaging set up

TIRF-DNA-PAINT was performed on a custom Nikon Eclipse Ti-2 microscope equipped with a 100×1.49 NA TIRF objective, 560 nm laser (MPB Communications), flat-top beam shaper (piShaper 6_6_VIS, AdlOptica), and Hamamatsu ORCA-Fusion BT sCMOS camera (130 nm/pixel after 2×2 binning). Acquisitions comprised 40,000 frames at 100 ms integration time. Raw data were processed in Picasso11 with redundant cross-correlation drift correction using gold nanoparticles as fiducials. Individual Env trimers were identified by co-localisation with Gag–mTurquoise2. **Quantitative DNA-PAINT analysis (qPAINT)** Gp120 copy numbers per virion were determined by qPAINT31, which infers protein copy number from the predictable binding kinetics of DNA imager and docking strands. Wild-type and EnvΔCT virions were compared; statistical differences were assessed using Prism 9.1 (GraphPad).

### DNA-PAINT TIRF acquisition and analysis

First, an image with a 488 nm CW laser was taken for 200 ms and a power density of 0.9 kW/cm2 to localize the position of the Gag-mAmetrine center of the virus particles on the previously described custom built microscope. Subsequent images were taken as a sequence of 100,000 frames, using an integration time of 100 ms and a density power of 2.3 kW/cm2.

The raw fluorescence videos were processed for super-resolution reconstruction using the Picasso software package11. Drift correction was applied using redundant cross-correlation, utilizing gold particles as fiducial markers for cellular experiments. Typically, around 50 nanoparticles were detected within the field of view, providing reliable drift correction, particularly for lateral shifts at the focal plane (z = 0).

Image rendering was carried out using the Render module of Picasso. Each localization was represented as a Gaussian spot, with the spread corresponding to the individual localization precision. For each pixel, the intensities of overlapping Gaussian spots were summed to determine the pixel intensity, which was then visualized using an appropriate colormap. To differentiate viruses from nonspecific binding or endosomes, the rendered DNA-PAINT image was aligned with the diffraction-limited image of the Gag channel using the gold nanoparticles. Only structures with the Gag signal were analyzed.

Protomer distance distributions were fitted with a two-component Gaussian mixture model (GMM) to resolve distinct conformational populations. Only the maximal inter-protomer distance (L*max*) was analysed to mitigate 3D-to-2D projection effects (Supplementary Figs. 11,12). Parameters (μ_1_, σ_1_, ω_1_, μ_2_, σ_2_, ω_2_) were estimated by direct maximum-likelihood optimisation, minimising the negative log-likelihood of the data. Parameter uncertainties were estimated from the diagonal of the inverse Hessian matrix returned by the optimiser at convergence, providing asymptotic standard errors on the fitted weights, means, and standard deviations. Fits were performed independently for mature, immature, mature+CD4, and immature+CD4 conditions. Localisation precision was determined by CRLB metrics.

### 3D-DNA-PAINT simulations

The simulations of the trimers shown in Figure S11 were performed using the Simulate package of Picasso software^37^. Parameters such as kinetic rates, imager strand concentration, excitation power density, background level, camera integration time, pixel size, and total number of frames were set to match the experimental conditions, ensuring comparable localization precision and NeNA^38^ values.

### Statistical analysis

Data analysis, statistical comparisons, and graphing were performed with Prism v9.1 (GraphPad Software) and custom-made python codes. All experiments were performed in at least three independent biological replicates unless stated otherwise. For the comparisons shown in Figure 2F, differences in the proportion of open, intermediate, and closed Env populations were assessed using two complementary approaches. Within-condition comparisons between conformational states (e.g., open vs intermediate, open vs closed) were assessed using a χ² test, treating the states as mutually exclusive subpopulations of the same set of virions. Across-condition comparisons of the same state (e.g., closed in mature vs closed in immature) were instead assessed using a two-proportion z-test, appropriate for comparing a single proportion measured in two independent samples. QPAINT gp120 per virus values for mature, immature, and ΔCT Env (Figure 3C) were compared using a Kruskal-Wallis test with Dunn’s post-hoc correction for multiple comparisons (Mature vs immature, mature vs ΔCT, Immature vs ΔCT).

### Modelling of Env–NbALFA and Env–NbALFA–CD4 structures

Complexes of Env–NbALFA and Env–NbALFA–CD4 were modelled using the amino acid sequences of our constructs and the AlphaFold 3 (AF3) web server. Asparagine-linked glycosylation sites on Env and CD4 were predicted using the NetNGlyc 1.0 web server and modelled as branched glycans using AF3 (code NAG(NAG(MAN(MAN(MAN)(MAN(MAN)(MAN)))))). The amino acid sequence of the anti-ALFA-tag nanobody from PDB ID 6I2G (ref. 3) was used for modelling. Three binary AF3 Env– NbALFA models or three ternary Env–NbALFA–CD4 models were superposed onto Env trimers representing closed (PDB ID 5T3X, ref. 4) and open (PDB ID 5VN3, ref. 5) conformations. The α-helix of Env (Met100–Leu116), proximal to the threefold symmetry axis and resolved in both conformations, was used for superposition. Distances between Cα atoms of NbALFA Val122 were measured, and structures were visualised using the PyMOL Molecular Graphics System, v. 3.0 (Schrödinger, LLC). N- and C-termini of Env (Met1–Lys32 and Lys496–Leu850) and glycans are not shown, and only the first two domains of CD4 are shown, for simplicity.

