## Supplementary Information for "Gag proteolysis drives HIV-1 envelope conformational closure through the gp41 cytoplasmic tail"

### Supplementary Figures

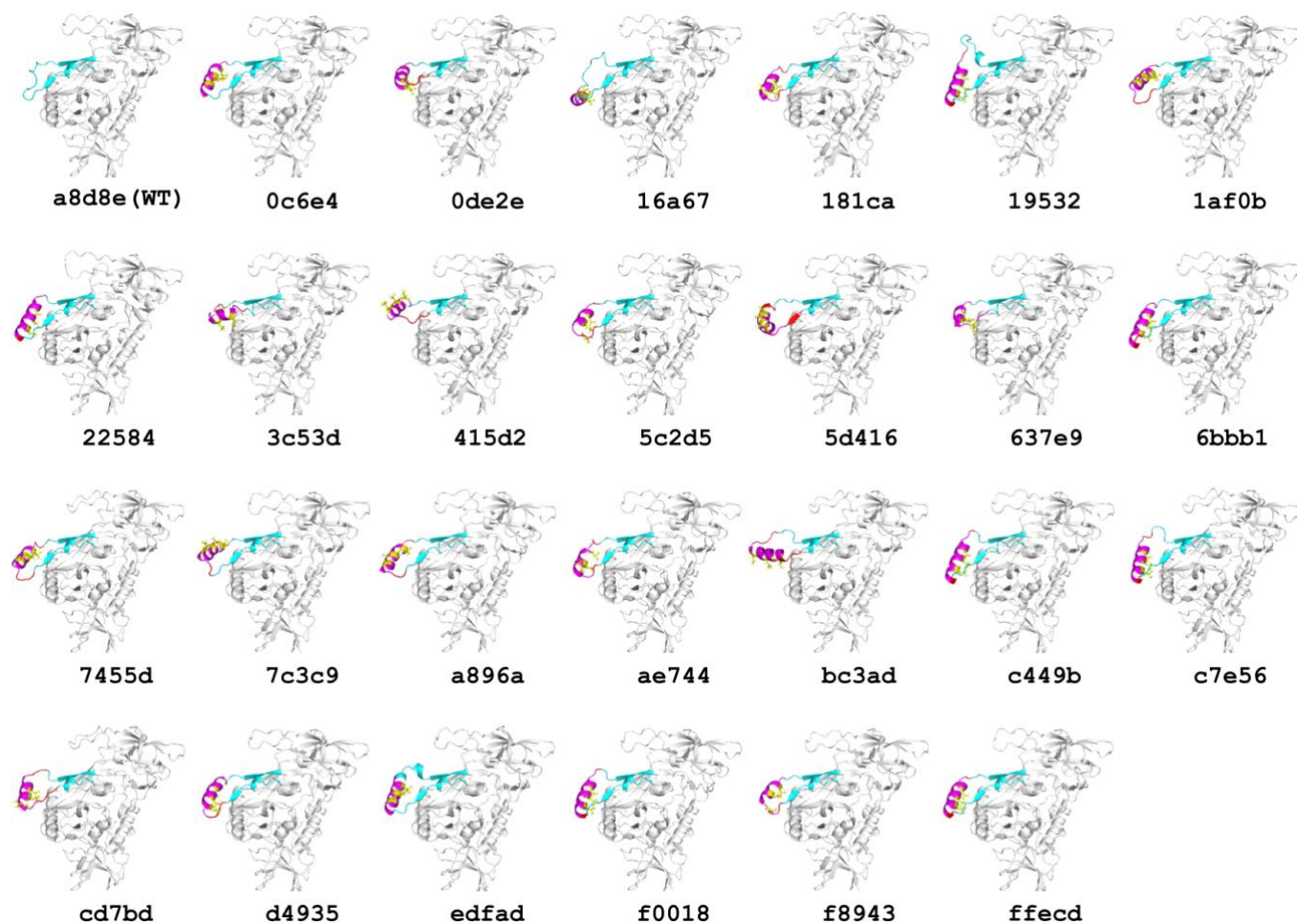

**Supplementary Figure 1. Overview of JR-FL wildtype (WT) and V4-ALFA-tag labelled gp120 structures.** Alphafold2 structure predictions of the JR-FL Env gp120 with ALFA-tag insertions across the variable loop 4 (V4). The highest-ranked model for each candidate insertion site (UID labels indicated) is shown. The ALFA-tag is highlighted in magenta, the V4 loop in cyan, and nanobody-binding pocket leucines in yellow. The wild-type (WT) JR-FL structure is included for comparison.

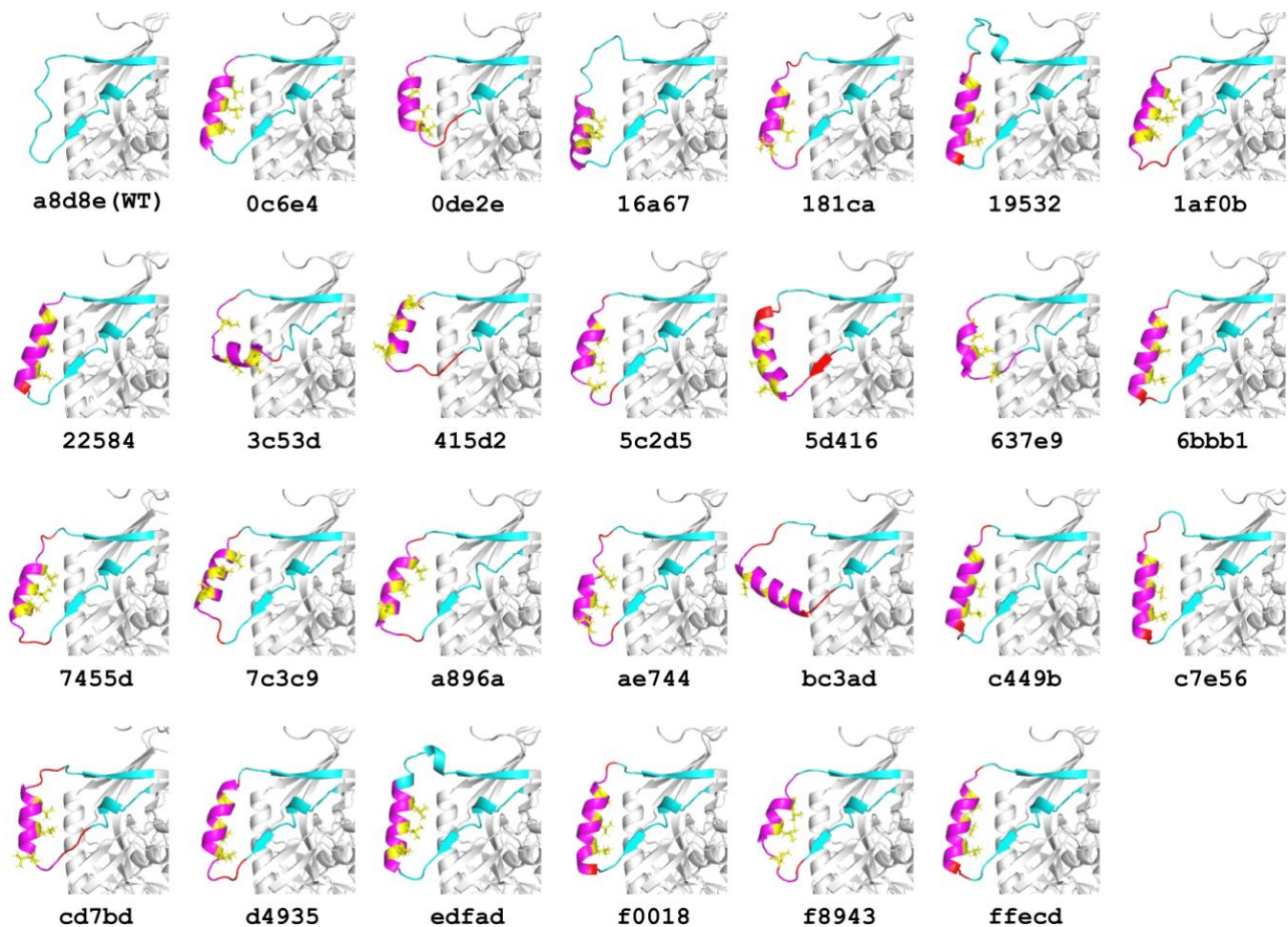

**Supplementary Figure 2. Zoomed, rotated view of the V4 loop region of the WT and labelled gp120 structures from Supplementary Figure 1.** Alphafold2 structure predictions (and UID labels) of the JR-FL Env gp120 with ALFA-tag insertions across the variable loop 4 (V4). Colouring as in Supplementary Figure 1.

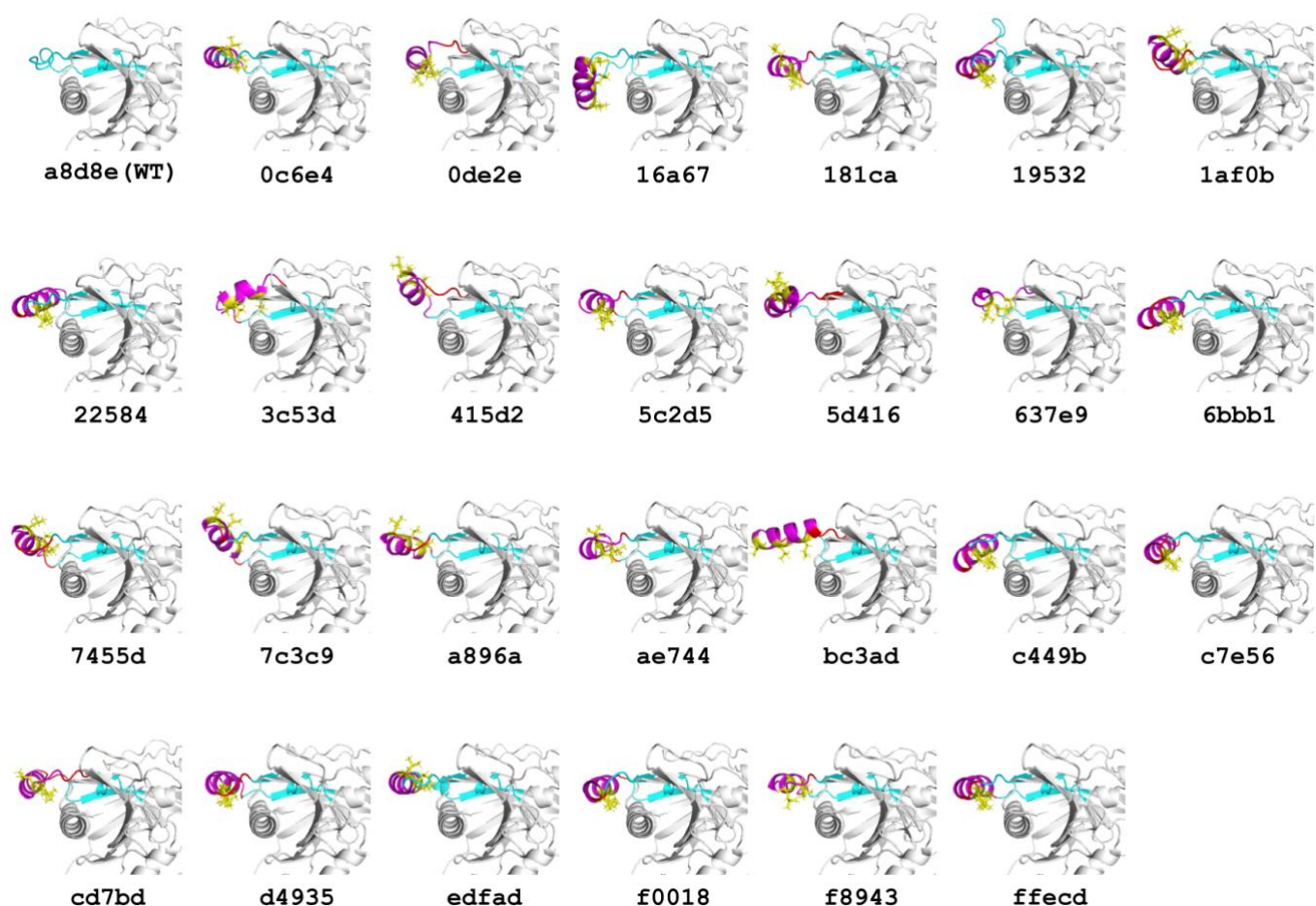

**Supplementary Figure 3. Zoomed, alternative rotated view of the V4 loop region of the WT and labelled gp120 structures from Supplementary Figure 1.** Alphafold2 structure predictions (and UID labels) of the JR-FL Env gp120 with ALFA-tag insertions across the variable loop 4 (V4), providing additional perspectives on tag conformation and pocket leucine orientation. Colouring as in Supplementary Figure 1.

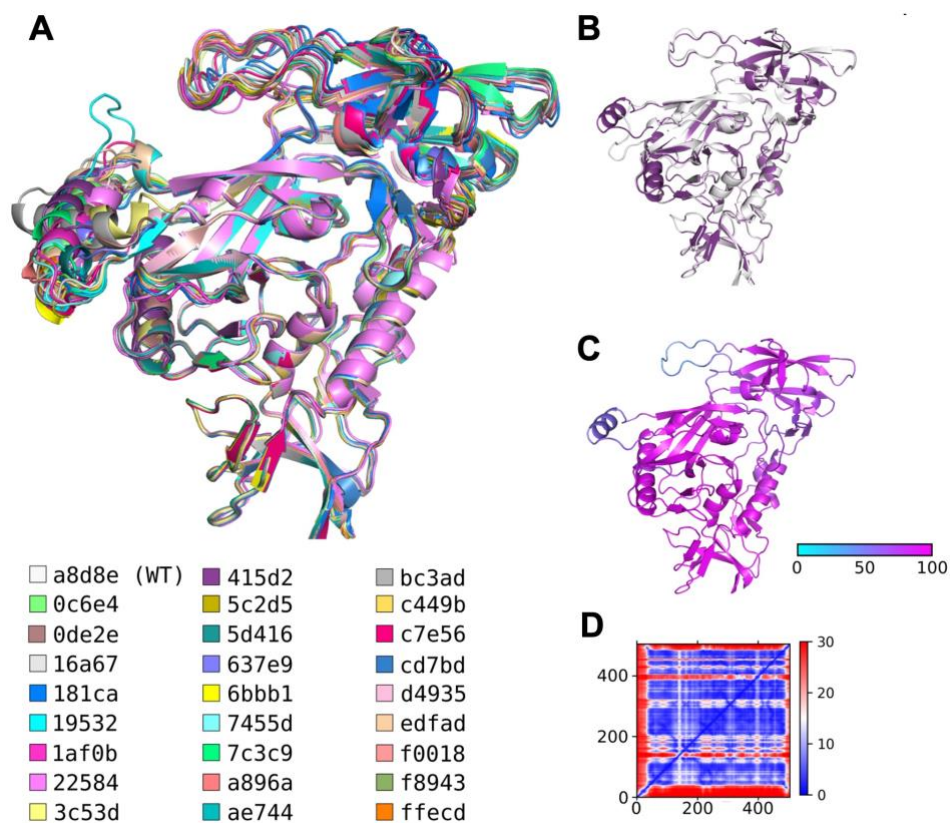

**Supplementary Figure 4. Structural alignment and quality metrics for ALFA-tag insertion proposals.** (A) All highest-ranked insertion models aligned to the JR-FL wild-type structure. (B) Model UID 415d2 (purple) aligned to JR-FL WT (grey), highlighting minimal backbone perturbation. (C) Model UID 415d2 colored by predicted Local Distance Difference Test (pLDDT) score, indicating per-residue confidence. (D) Predicted Aligned Error (PAE) heatmaps for 415d2, reporting inter-residue distance uncertainty.

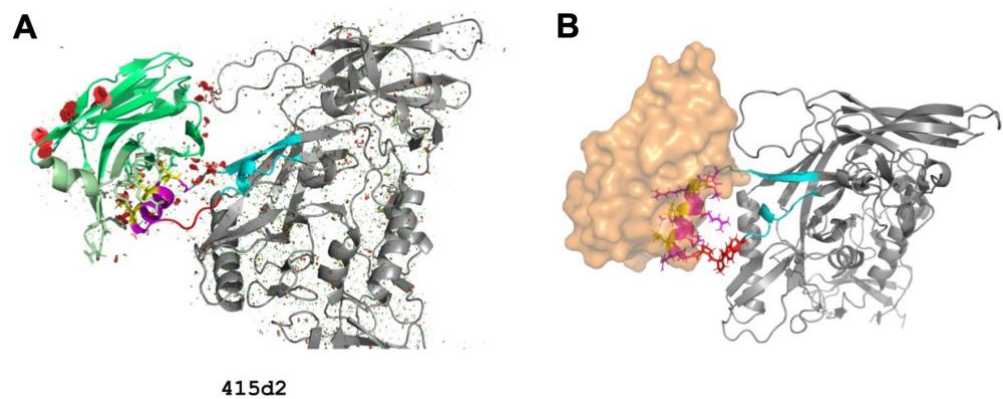

**Supplementary Figure 5. Nanobody docking onto the 415d2 ALFA-tag insertion model. (A, B)** JRFL V4-ALFA-tag UID 415d2 with the inserted tag aligned to the ALFA-tag from PDB ID 6IG2 to indicate where the NbALFA (shown in green cartoon or orange surface renderings) may bind. Steric clashes are indicated with red polygons. Nanobody docking onto the 415d2 ALFA-tag insertion model.

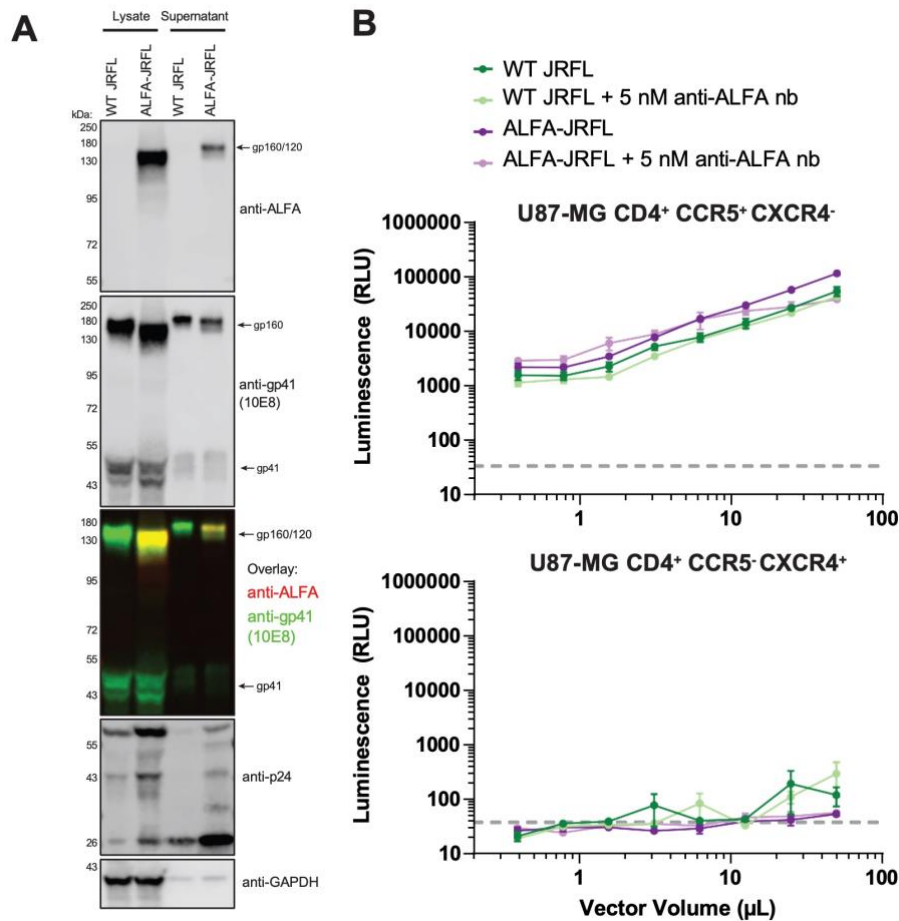

**Supplementary Figure 6. Biochemical and functional validation of ALFA-tagged JR-FL Env. (A)** Western blot of HEK293T cell lysates and concentrated supernatant from cells transfected to produce HIV-1-based lentivirus pseudo typed with either wild-type or ALFA-tagged (UID 415d2) JR-FL Env, probed for gp120 and gp41 to confirm correct processing and incorporation. **(B)** Single-cycle infectivity of WT and 415d2 pseudovirus on U87-MG CD4<sup>+</sup> cells expressing CCR5 or CXCR4, with or without a pre-incubation with 5 nM nbALFA (30 min). Infection is quantified as relative luminescence units. ALFA-tag insertion and nbALFA binding do not impair viral entry.

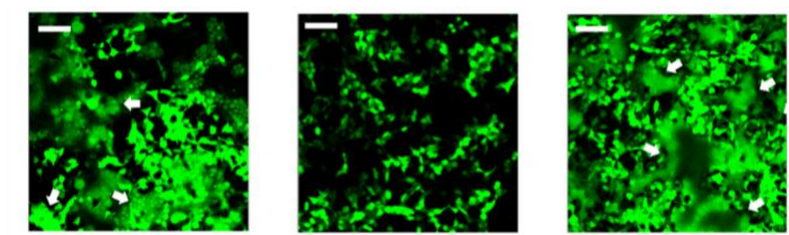

**Supplementary Figure 7. Cell-cell fusion assays for JR-FL V4-ALFA-tag UID 415d2.** Permissive TZM-bl target cells were exposed to HEK T293 effector cells expressing: wildtype JR-FL Env (i.e, no V4 label, left), empty vector (middle), or JR-FL V4-ALFA-tag UID 415d12 (right). Syncytia (white arrows) indicates productive fusion. Scale bar, 10 μm. Fusion activity is retained in the ALFA-tagged construct.

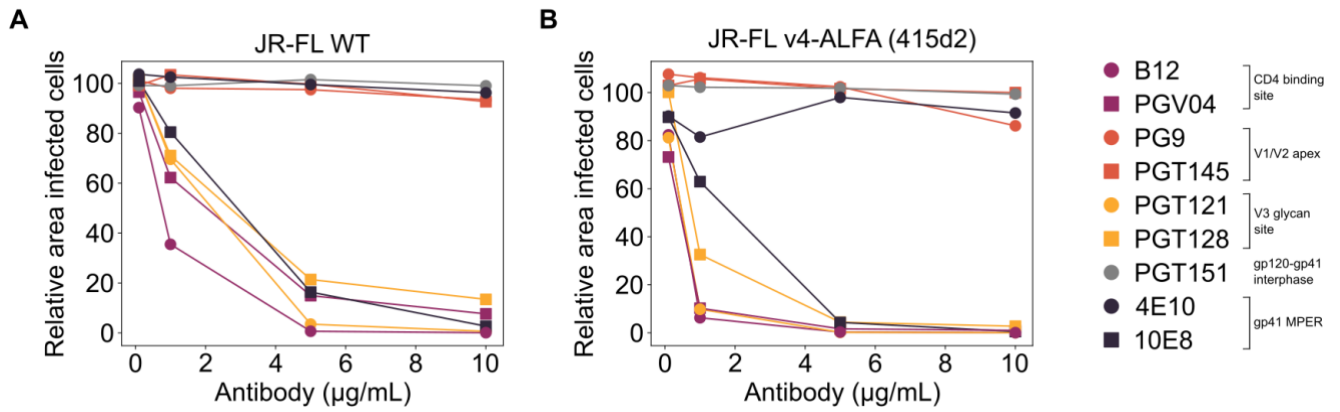

**Supplementary Figure 8. Infectivity-neutralization assay in-vitro testing broadly neutralizing antibodies (bnAbs) targeting different epitopes of HIV-1 Env.** Increasing concentrations of bnAbs targeting distinct Env epitopes — CD4 binding site (b12, PGV04), V1/V2 apex (PG9, PGT145), V3 loop (PGT121, PGT128), gp120–gp41 interface (PGT151), and gp41 MPER (4E10, 10E8) — were applied to TZM-bl cells infected with equal amounts of HIV-1 viruses JR-FL WT (**A**) and decorated with JR-FL Env V4-ALFA-tag (**B**). Under these conditions, b12 showed the most potent neutralisation and was selected for subsequent conformation-sensitive binding experiments, consistent with its known selectivity for open Env conformations.

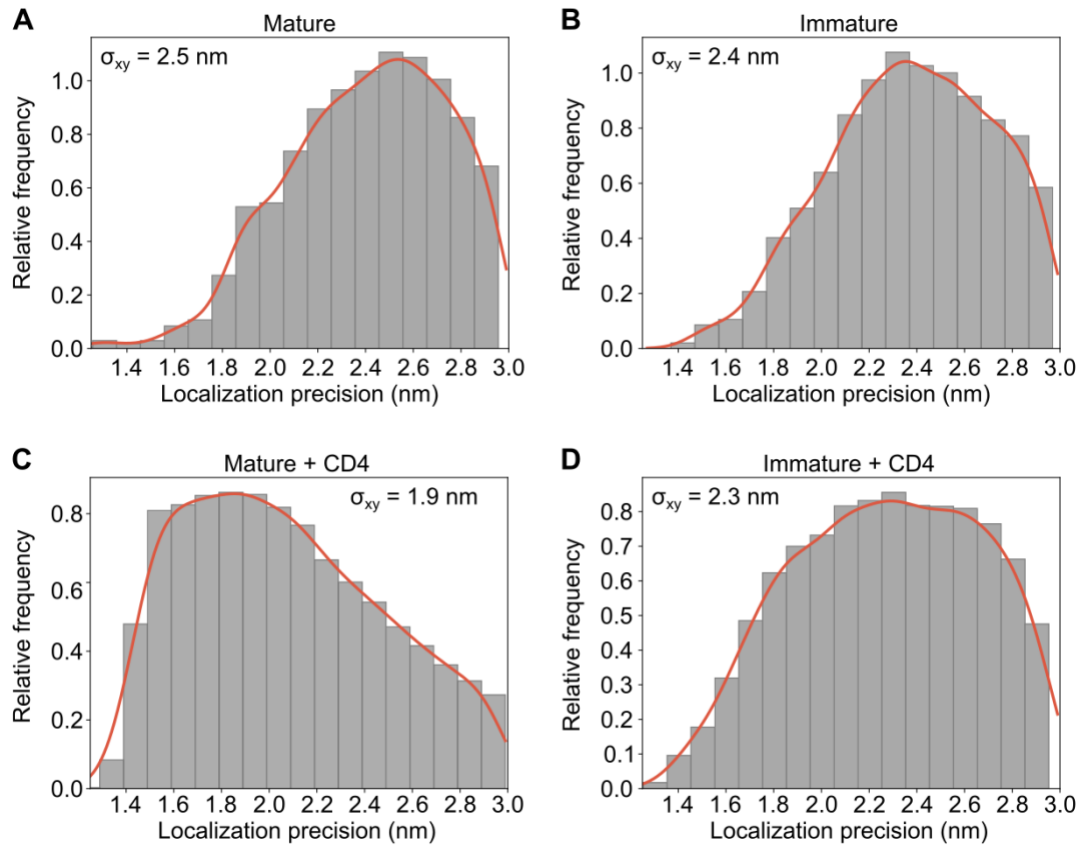

**Supplementary Figure 9. Localization precision distributions for DNA-PAINT imaging.**

Distributions of localization precision ($\sigma_{xy}$) for all super-resolution DNA-PAINT datasets: **(A)** mature virions, **(B)** immature virions, **(C)** mature virions treated with soluble CD4, and **(D)** immature virions treated with soluble CD4. Values reported are the mode of each distribution.

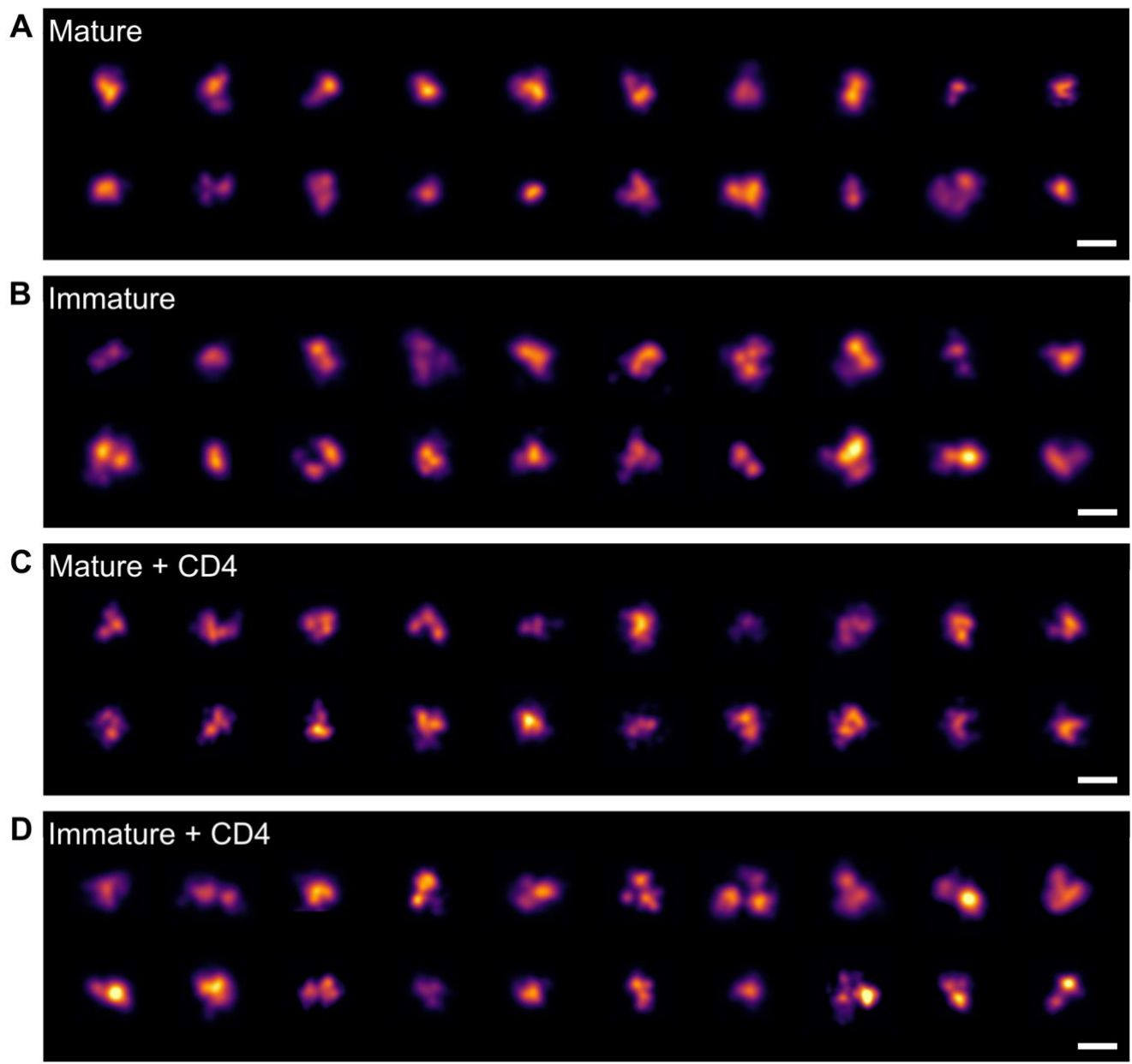

**Supplementary Figure 10. Representative DNA-PAINT images of individual Env trimers across all conditions.** Single-trimer images from across the field of view for (A) mature, (B) immature, (C) mature + sCD4, and (D) immature + sCD4 virions, illustrating the range of trimer morphologies observed under each condition. Scale bar: 20 nm.

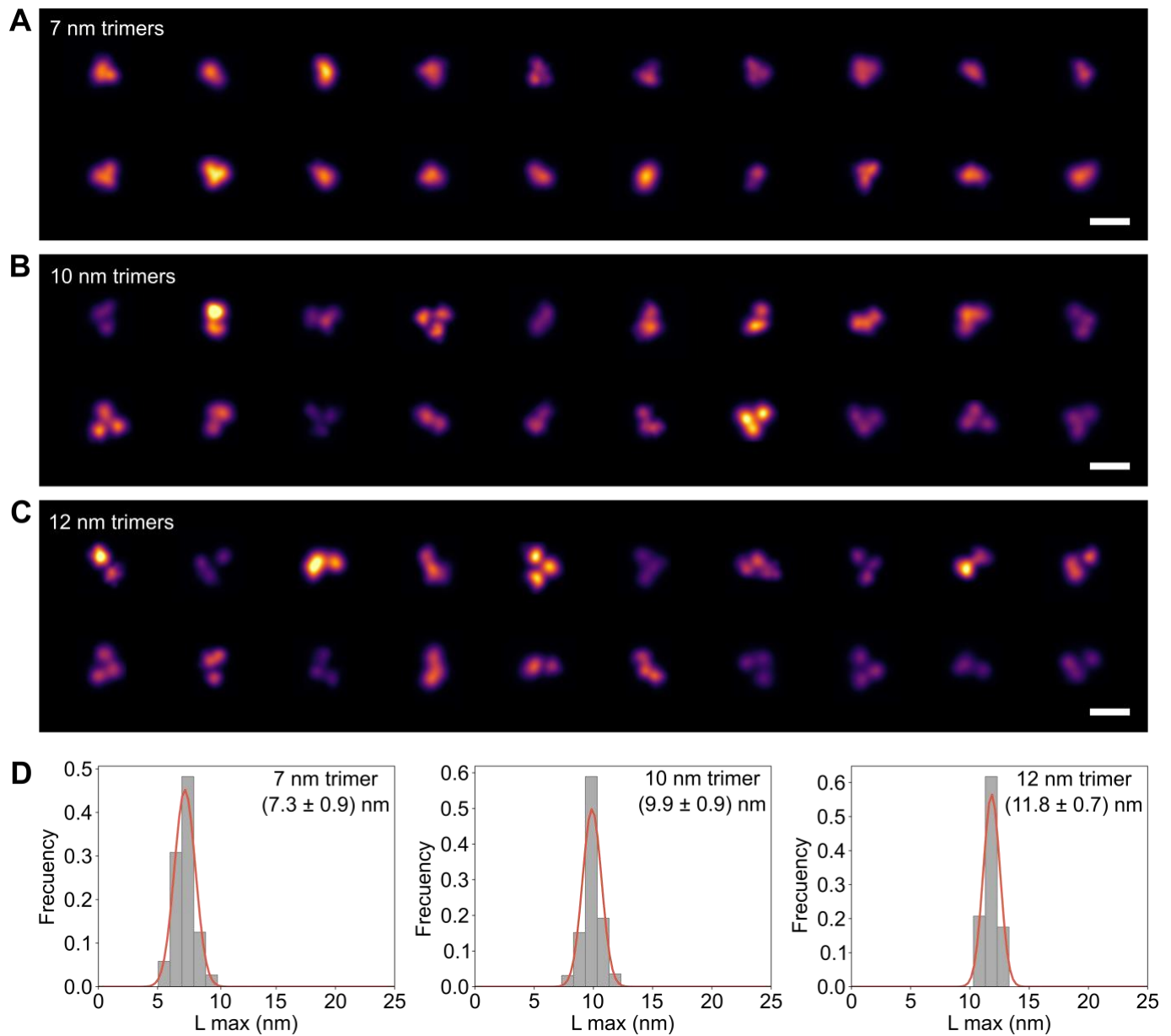

**Supplementary Figure 11: Simulated DNA-PAINT 2D projections of 3D equilateral trimers at varied inter-protomer distances.** 2D spatial projections derived from simulations of 3D equilateral trimer geometries. Panels illustrate the resulting 2D localization renders for (A) 7 nm, (B) 10 nm, (C) and 12 nm inter-protomer distance. Scale bar: 20 nm. (D) Distributions of L max for each inter-protomer distance fitted with a three-component Gaussian mixture model. The mean and standard deviation for each fitted Gaussian component are provided inset for each distribution.

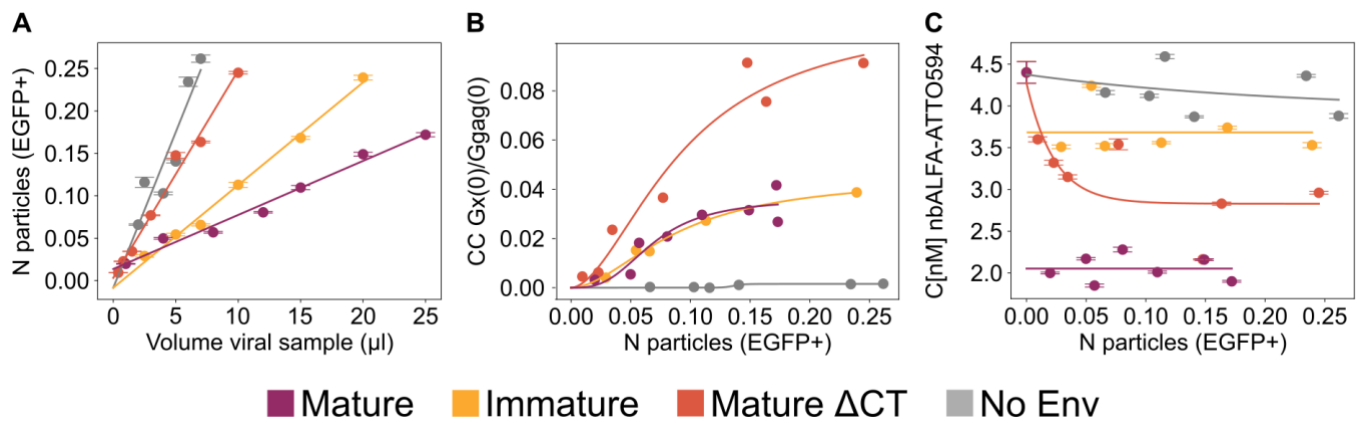

**Supplementary Figure 12. Two-color FLCS virometry of JR-FL Env V4-ALFA-tag UID 415d2.** (A) Number of viruses as a function of volume for HIV-1 viruses lacking Env (No Env, grey), bearing JR-FL Env V4-ALFA with the C terminus of gp41 truncated (ΔCT) (orange), or bearing full-length JR-FL V4-ALFA in immature (yellow) or mature (purple) form. All particles carry Gag-EGFP as a virion reporter and are decorated with nbALFA-ATTO594. (B) Cross-correlation curves free from artifacts coming from two colour FLCS for the conditions in (A). (C) Concentration of nbALFA-ATTO594 plotted against viral particle number (Gag-EGFP/WT Gag ratio 10:90).

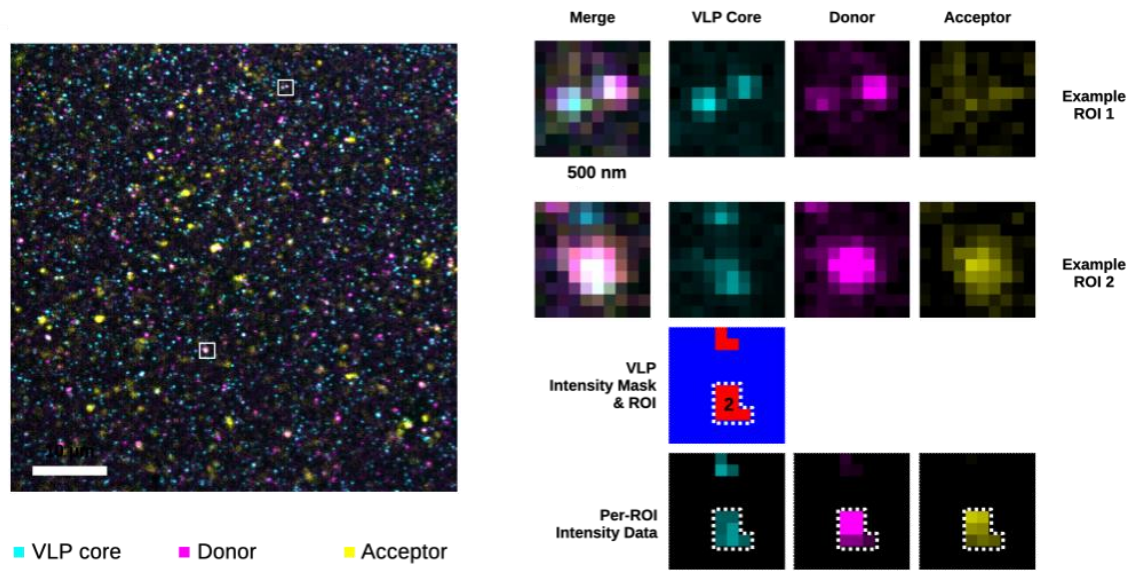

**Supplementary Figure 13. FRET imaging configuration.** Micrograph showing HIV-1 virions labelled with Gag-mAmetrine (VLP core, cyan), decorated with JR-FL-V4-ALFA-tag (Atto488, Donor in Red and Atto594 Acceptor in Yellow). Scale bar 2 μm. The white squares are shown on the right-hand side, and both the merged colors and the VLP core, the Donor and the Acceptor signal for FRET calculation are shown. Only particles with positive VLP core were taken into consideration for FRET Env calculation. An intensity mask and ROI was used as a mask to recover the photons from the three channels and the FRET calculation was performed with the Donor and Acceptor channels with at least 100 photons per channel as specified in material and methods.

| UID | V4 ... | Padding | ALFA-Tag | Padding | ... V4 |
| --- | --- | --- | --- | --- | --- |
| 0c6e4 | CNSTQLFNSTWNNN |  | <u>SRLEEELRRRLTE</u> |  | GNTITLPC |
| 0de2e | CNSTQLFN | PP | <u>SRLEEELRRRLTE</u> |  | GNTITLPC |
| 16a67 | CNSTQLFNSTWNNN |  | <u>SRLEEELRRRLTE</u> | TEGSNNTEGNTITLPC |  |
| 181ca | CNSTQLFNSTW | A | <u>SRLEEELRRRLTE</u> | PP | GNTITLPC |
| 19532 | CNSTQLFNSTWNNN | P | <u>SRLEEELRRRLTE</u> | P | TEGSNNTEGNTITLPC |
| 1af0b | CNSTQLFNSTW | PPPA | <u>SRLEEELRRRLTE</u> | A | GNTITLPC |
| 22584 | CNSTQLFNSTWNNN | A | <u>SRLEEELRRRLTE</u> |  | GNTITLPC |
| 3c53d | CNSTQLFN | P | <u>SRLEEELRRRLTE</u> | P | GNTITLPC |
| 415d2 * | CNSTQLFN | FFPP | <u>SRLEEELRRRLTE</u> |  | GNTITLPC |
| 5c2d5 | CNSTQLFNSTW | P | <u>SRLEEELRRRLTE</u> | P | GNTITLPC |
| 5d416 | CNSTQLFN | SRL | <u>SRLEEELRRRLTE</u> | RRL | TEGNTITLPC |
| 637e9 * | CNSTQLFN |  | <u>SRLEEELRRRLTE</u> |  | GNTITLPC |
| 6bbb1 | CNSTQLFNSTWNN | PPP | <u>SRLEEELRRRLTE</u> | A | GNTITLPC |
| 7455d | CNSTQLFNSTWN | PPP | <u>SRLEEELRRRLTE</u> | PP | GNTITLPC |
| 7c3c9 | CNSTQLFNSTWNN | PP | <u>SRLEEELRRRLTE</u> | P | GNTITLPC |
| a896a | CNSTQLFNSTWN | PP | <u>SRLEEELRRRLTE</u> | A | GNTITLPC |
| ae744 | CNSTQLFNSTW | P | <u>SRLEEELRRRLTE</u> | PP | GNTITLPC |
| bc3ad | CNSTQLFN | FFP | <u>SRLEEELRRRLTE</u> | PFF | GNTITLPC |
| c449b | CNSTQLFNSTWNNN | PP | <u>SRLEEELRRRLTE</u> | P | GNTITLPC |
| c7e56 | CNSTQLFNSTWNN | PA | <u>SRLEEELRRRLTE</u> | AP | TEGNTITLPC |
| cd7bd | CNSTQLFN | TQL | <u>SRLEEELRRRLTE</u> | TQL | GNTITLPC |
| d4935 | CNSTQLFNSTW | PPP | <u>SRLEEELRRRLTE</u> | A | GNTITLPC |
| edfad | CNSTQLFNSTWNNN |  | <u>SRLEEELRRRLTE</u> | GSNNTEGNTITLPC |  |
| f0018 | CNSTQLFNSTWNN | P | <u>SRLEEELRRRLTE</u> | P | GNTITLPC |
| f8943 | CNSTQLFNSTW | P | <u>SRLEEELRRRLTE</u> |  | GNTITLPC |
| ffecd | CNSTQLFNSTWNN | FP | <u>SRLEEELRRRLTE</u> | P | GNTITLPC |
| a8d8e (JR-FL WT) | CNSTQLFNSTWNNNTEGSNNTEGNTITLPC |  |  |  |  |

\* Expressed and tested

<u>TE</u> = ALFA-tag & V4 homology

**Supplementary Table 1. List of ALFA-tag insertion proposals.** A panel of ALFA-tag insertion variants was designed by introducing the ALFA-tag sequence (SRLEEELRRRLTE) at different positions within the V4 loop of JR-FL Env. Flanking residues ("Padding") were included to preserve local sequence context and structural compatibility. The wild-type (WT) V4 sequence is shown for reference (UID a8d8e). Variants marked with an asterisk (\*) were successfully expressed and selected for experimental characterization.

| Section | Comparison | State 1 | State 2 | Test | p-value | Significance |
| --- | --- | --- | --- | --- | --- | --- |
| 1. Within condition | Mature | Open | Intermediate | $\chi^2$ | 5.34E-01 | ns |
| 1. Within condition | Mature | Open | Closed | $\chi^2$ | 5.21E-02 | ns |
| 1. Within condition | Mature | Intermediate | Closed | $\chi^2$ | 1.06E-02 | * |
| 1. Within condition | Immature | Open | Intermediate | $\chi^2$ | 1.91E-09 | *** |
| 1. Within condition | Immature | Open | Closed | $\chi^2$ | 1.64E-26 | *** |
| 1. Within condition | Immature | Intermediate | Closed | $\chi^2$ | 1.43E-07 | *** |
| 1. Within condition | Mature+CD4 | Open | Intermediate | $\chi^2$ | 6.27E-01 | ns |
| 1. Within condition | Mature+CD4 | Open | Closed | $\chi^2$ | 4.74E-16 | *** |
| 1. Within condition | Mature+CD4 | Intermediate | Closed | $\chi^2$ | 1.24E-17 | *** |
| 1. Within condition | Immature+CD4 | Open | Intermediate | $\chi^2$ | 1.35E-15 | *** |
| 1. Within condition | Immature+CD4 | Open | Closed | $\chi^2$ | 8.94E-48 | *** |
| 1. Within condition | Immature+CD4 | Intermediate | Closed | $\chi^2$ | 3.59E-13 | *** |
| 2. Across conditions | Mature vs Immature | Closed | - | z-test | 1.70E-12 | *** |
| 2. Across conditions | Mature vs Mature+CD4 | Closed | - | z-test | 6.32E-13 | *** |
| 2. Across conditions | Mature vs Immature+CD4 | Closed | - | z-test | 6.46E-17 | *** |
| 2. Across conditions | Immature vs Mature+CD4 | Closed | - | z-test | 9.06E-01 | ns |
| 2. Across conditions | Immature vs Immature+CD4 | Closed | - | z-test | 6.70E-01 | ns |
| 2. Across conditions | Mature+CD4 vs Immature+CD4 | Closed | - | z-test | 7.61E-01 | ns |
| 2. Across conditions | Mature vs Immature | Intermediate | - | z-test | 5.61E-01 | ns |
| 2. Across conditions | Mature vs Mature+CD4 | Intermediate | - | z-test | 3.17E-04 | *** |
| 2. Across conditions | Mature vs Immature+CD4 | Intermediate | - | z-test | 4.33E-01 | ns |
| 2. Across conditions | Immature vs Mature+CD4 | Intermediate | - | z-test | 8.04E-04 | *** |
| 2. Across conditions | Immature vs Immature+CD4 | Intermediate | - | z-test | 8.81E-01 | ns |
| 2. Across conditions | Mature+CD4 vs Immature+CD4 | Intermediate | - | z-test | 1.28E-04 | *** |
| 2. Across conditions | Mature vs Immature | Open | - | z-test | 7.08E-08 | *** |
| 2. Across conditions | Mature vs Mature+CD4 | Open | - | z-test | 1.16E-02 | * |
| 2. Across conditions | Mature vs Immature+CD4 | Open | - | z-test | 9.50E-09 | *** |
| 2. Across conditions | Immature vs Mature+CD4 | Open | - | z-test | 8.45E-04 | *** |
| 2. Across conditions | Immature vs Immature+CD4 | Open | - | z-test | 6.92E-01 | ns |
| 2. Across conditions | Mature+CD4 vs Immature+CD4 | Open | - | z-test | 4.41E-04 | *** |

**Supplementary Table 2.** The proportions of closed, intermediate, and open conformational states were estimated by fitting a two-component Gaussian mixture model (GMM) to the $L_{\max}$ distance measured for each Env trimer. Thresholds of 8 nm and 11 nm were selected to delineate closed, intermediate, and open populations, based on their position relative to the closed- and open-state component means obtained from Gaussian mixture fitting across all four conditions (closed-component means: 7.1–10.0 nm; open-component means: 11.9–16.0 nm). These values fall within the gap separating the two fitted components for every condition and were not adjusted to favour any single dataset. Sensitivity analysis ($\pm 1$ nm on each boundary) confirmed that classification into closed and open states, and the corresponding statistical comparisons between mature and immature/CD4-bound Env, were unchanged across this range. Differences involving the intermediate population were more sensitive to exact threshold placement, as expected for a bin bounded on both sides, and are therefore reported as descriptive rather than a basis for statistical claims. Comparisons among states within the same condition were assessed using a chi-squared test based on state-assigned bin counts. Comparisons of individual states across conditions were performed using a two-sample z-test applied to the GMM-derived mixing weights and their associated standard errors. Statistical significance is indicated as follows: \*$p < 0.05$, \*\*$p < 0.01$, \*\*\*$p < 0.001$, ns: not significant
